# Transcription bodies regulate gene expression by sequestering CDK9

**DOI:** 10.1101/2022.11.21.517317

**Authors:** Martino Ugolini, Ksenia Kuznetsova, Haruka Oda, Hiroshi Kimura, Nadine L. Vastenhouw

## Abstract

The localization of transcriptional activity in specialized transcription bodies is a hallmark of gene expression in eukaryotic cells. It remains unclear, however, if and how they affect gene expression. Here, we disrupted the formation of two prominent endogenous transcription bodies that mark the onset of zygotic transcription in zebrafish embryos and analysed the effect on gene expression using enriched SLAM-Seq and live-cell imaging. We find that the disruption of transcription bodies results in downregulation of hundreds of genes, providing experimental support for a model in which transcription bodies increase the efficiency of transcription. We also find that a significant number of genes are upregulated, counter to the suggested stimulatory effect of transcription bodies. These upregulated genes have accessible chromatin and are poised to be transcribed in the presence of the two transcription bodies, but they do not go into elongation. Live-cell imaging shows that the disruption of the two large transcription bodies enables these poised genes to be transcribed in ectopic transcription bodies, suggesting that the large transcription bodies sequester a pause release factor. Supporting this hypothesis, we find that CDK9, the kinase that releases paused polymerase II, is highly enriched in the two large transcription bodies. Importantly, overexpression of CDK9 in wild type embryos results in the formation of ectopic transcription bodies and thus phenocopies the removal of the two large transcription bodies. Taken together, our results show that transcription bodies regulate transcription genome-wide: the accumulation of transcriptional machinery creates a favourable environment for transcription locally, while depriving genes elsewhere in the nucleus from the same machinery.

---

RNA polymerase II (RNAPII) and transcription factors are often concentrated in specialized transcription bodies ^1–10^, and it has been suggested that these can bring together multiple genes ^11–14^. This has led to the hypothesis that transcription bodies increase the efficiency of transcription by promoting the biomolecular interactions underlying gene expression ^15–17^. Analysis of gene expression in artificially induced condensates supports this model ^18^. To understand how transcription bodies affect gene expression *in vivo*, however, it is necessary to analyse endogenous transcription bodies. This has been difficult because transcription bodies are often small, short-lived, and numerous. Here, we use the onset of transcription in zebrafish embryos to overcome this problem. In zebrafish embryos, development is initially driven by maternally loaded RNA and protein (Fig. 1a) ^19^. Transcription gradually begins during a process called zygotic genome activation (ZGA). Before transcriptional activity can be seen throughout the nucleus, it is confined to two micron-sized transcription bodies (Fig. 1b) ^9,20–23^, nucleated by the *mir430* locus, which contains multiple copies of the *mir430* gene ^21,24^. These bodies are enriched for RNAPII Serine 5 and Serine 2 phosphorylation (Ser5P, Ser2P), which mark the initiating and elongating form of RNAPII, respectively, as well as the transcription factors Nanog and Sox19b ^9,20–22^.

**Figure 1.**
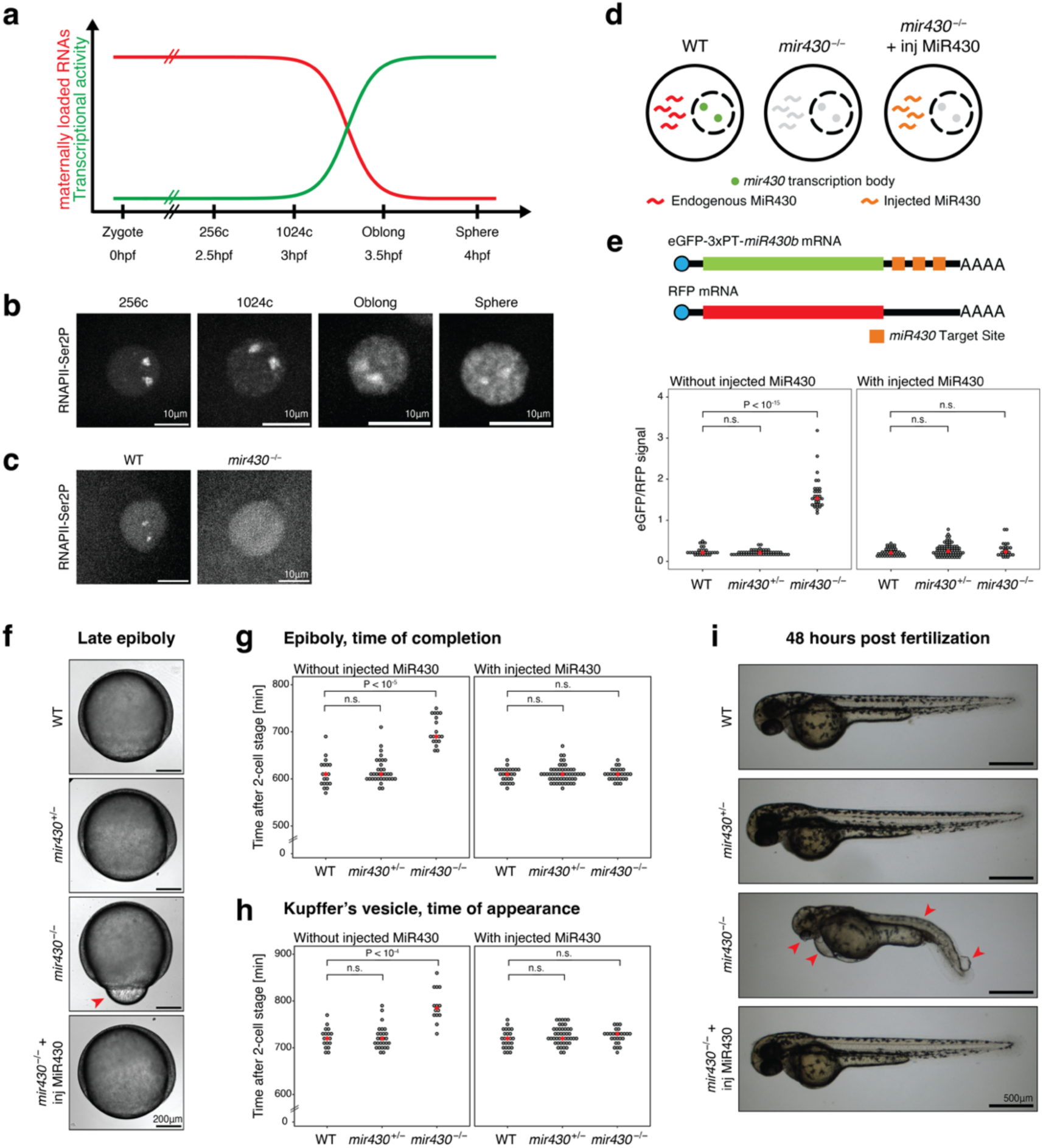
Disruption of *mir430* transcription bodies does not substantially impact development. **a.** Schematic representation of the maternal-to-zygotic transition in zebrafish embryos. Hpf, hours post fertilization. **b.** Visualization of elongating RNAPII (Ser2P) with antigen-binding fragments (Fabs) in WT embryos. Shown are representative micrographs of individual nuclei, extracted from a spinning disk confocal microscopy time lapse. **c.** Visualization of elongating RNAPII (Ser2P) with antigen-binding fragments (Fabs) in WT and *mir430*^-/-^ embryos at the 128-cell stage. Shown are representative micrographs of individual nuclei, extracted from a spinning disk confocal microscopy time lapse. **d.** Schematic representation of a nucleus in WT, *mir430*^-/-^ and *mir430*^-/-^ + inj MiR430 embryos. **e.** Rescue of *miR430* activity as assessed by comparing the expression of eGFP encoded by an mRNA with three perfect target sites for *miR430,* and RFP encoded by an mRNA without such sites, in different genotypes at 24hpf. Normalized eGFP signal in WT, *mir430*^+/-^ and *mir430*^-/-^ embryos without (left) and with (right) injected MiR430. N=3, 27≤n≤74. See Extended Data Fig. 1 for images. **f.** Rescue of *miR430* activity as assessed by epiboly progression. Shown are representative micrographs of embryos at late epiboly stage in different genotypes. The misregulation of yolk internalization in *mir430*^-/-^ embryos is indicated. **g.** Time at which epiboly is completed in different genotypes without (left), and with (right) injected MiR430 RNA. N=3, 19≤n≤53. **h.** Time at which Kupffer’s vesicle appears in different genotypes without (left), and with (right) injected MiR430. N=3, 14≤n≤44. **i.** Larvae at 48 hours post fertilization for different genotypes. Representative micrographs are shown. The malformation of trunk morphology and eye, the development of heart oedema, and the appearance of blisters at the tail tip in *mir430*^-/-^ embryos are indicated.

To investigate the role of transcription bodies in transcription, we generated a fish line in which the *mir430* locus is deleted ^9^, resulting in the specific disruption of the two *mir430* transcription bodies, as observed by the loss of Ser2P signal (Fig. 1c). *miR430* is required for downregulating maternally loaded transcripts, and its absence results in severe developmental phenotypes ^24,25^. Thus, effects on gene expression and development in the *mir430* mutant may be caused by the loss of the *mir430* transcription bodies, or the loss of the mature microRNA MiR430. To uncouple these two components and allow us to focus on the first, we supplemented mutant embryos with the mature form of microRNA MiR430 (Fig. 1d) ^24^, which restored degradation of mRNAs containing MiR430 binding sites (Fig. 1e and Extended Data Fig. 1). This rescued the delay in epiboly (Fig. 1f-g) and the aberrant expression pattern of *goosecoid* (Extended Data Fig. 2), which are characteristic for the loss of MiR430 ^24,25^. Embryos supplemented with mature microRNAs appeared normal, with the appearance of the Kupffer’s vesicle (Fig. 1h), and overall developmental progression (Fig. 1i and Extended Data Fig. 3) being indistinguishable from wild type (WT) and heterozygous siblings, and embryos developed into fertile fish. Thus, the disruption of two large transcription bodies does not result in obvious developmental defects, and we can use it to analyse the effect of transcription bodies on gene expression.

To assess the effect of the specific disruption of *mir430* transcription bodies on transcriptional activity, we developed eSLAM-Seq, which uses a combination of protocols to label, enrich, and detect nascent transcripts (Extended Data Fig. 4 and Methods), because the large amounts of maternally loaded RNA in early embryos (Fig. 1a) masks nascent RNAs in total RNA sequencing approaches. We used eSLAM-Seq to compare gene expression between WT and *mir430*^-/-^ + inj MiR430 embryos at the 256-cell stage, when transcriptional activity is largely restricted to the *mir430* transcription bodies in WT embryos (Fig. 1b). We observed a significant number of genes that were downregulated in the absence of the *mir430* transcription bodies (242) and a higher number of genes that were upregulated (716) (Fig. 2a-b and Extended Data Table 1). Plotting the genomic distribution of genes that were up- and downregulated showed that both classes of genes are distributed across the genome (Fig. 2c). We conclude that the absence of two large transcription bodies has a widespread effect on transcription.

**Figure 2.**
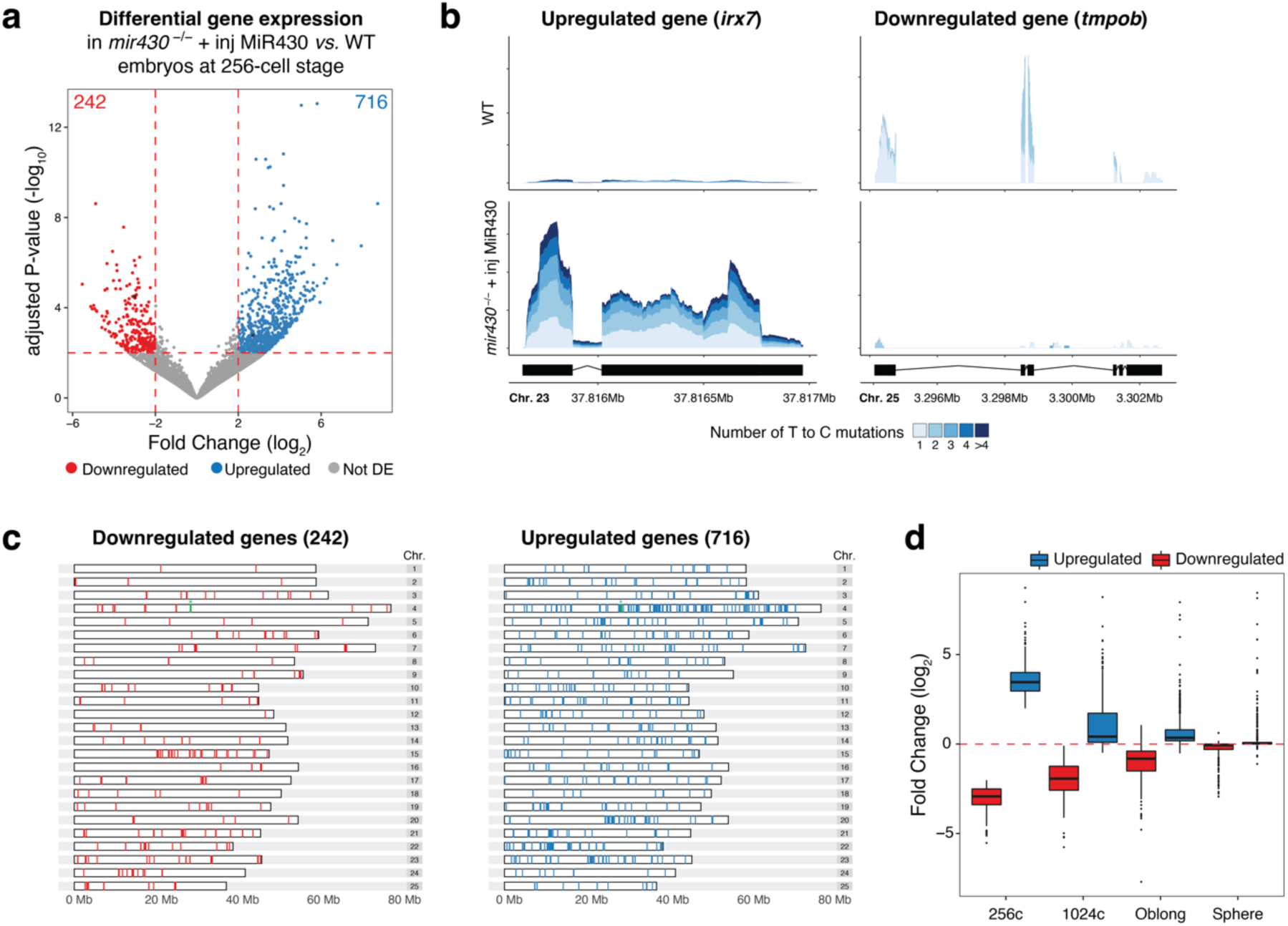
Disruption of *mir430* transcription bodies causes widespread misregulation of gene expression. **a.** Volcano plot showing up- and downregulated genes in *mir430*^-/-^ + inj MiR430 *vs.* WT embryos at 256-cell stage. Genes whose coverage is shown in Fig. 2b are shown in black. **b.** Coverage plot of *irx7* (upregulated) and *tmpob* (downregulated) in WT and *mir430*^-/-^ + inj MiR430. The single strata visualize the labelling degree of the reads. **c.** Distribution of up- and downregulated genes across the genome. The *mir430* locus on chromosome 4 is shown in green and labelled by an asterisk. **d.** Difference in average gene expression between *mir430*^-/-^ + inj MiR430 and WT embryos across stages for those genes that were identified to be differentially expressed at 256-cell stage.

**Figure 4.**
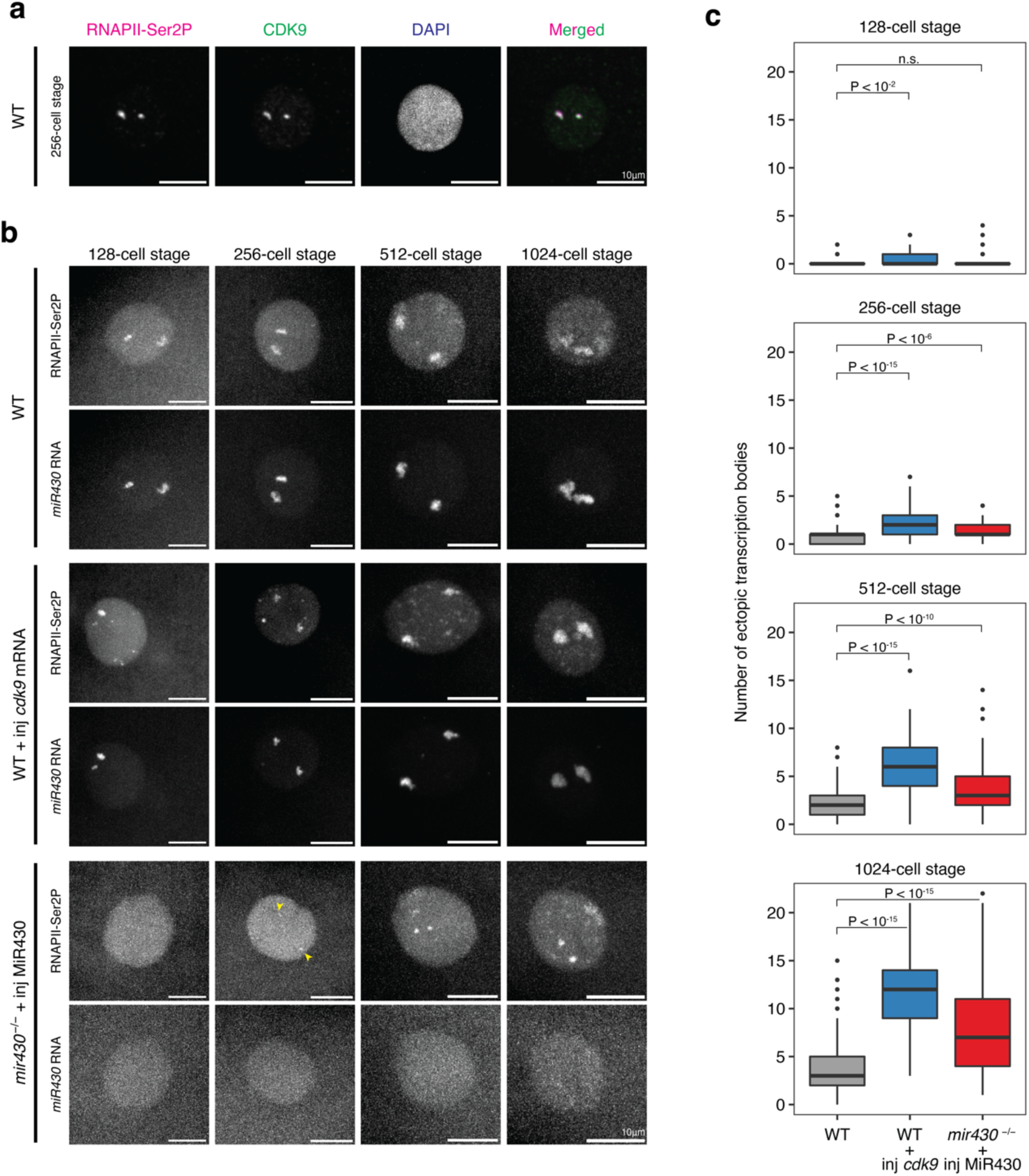
CDK9 sequestration in *mir430* transcription bodies inhibits transcription elongation elsewhere. **a.** Visualization of CDK9 and RNAPII-Ser2P by immunofluorescence in WT nuclei at 256c. Nuclei are labelled with DAPI. **b.** Visualization of RNAPII-Ser2P with antigen-binding fragments (Fabs), and *miR430* RNA with MOVIE in WT, WT + inj *cdk9* mRNA, and *mir430*^-/-^ + inj MiR430 embryos across stages. Shown are representative micrographs of individual nuclei, extracted from a spinning disk confocal microscopy time lapse. See Extended Data Fig. 11-14 for complete cell cycles. **c.** Quantification of ectopic transcription bodies as shown in b. N=3, 8≤n≤168. See Methods for a detailed description of quantification.

The identification of hundreds of up- and downregulated genes in *mir430*^-/-^ + inj MiR430 *vs.* WT embryos at the 256-cell stage contrasts with the observation that these embryos develop normally (Fig. 1). To reconcile these two observations, we generated eSLAM-Seq data for WT and *mir430*^-/-^ + inj MiR430 embryos at the 1024-cell, Oblong, and Sphere stage and compared it with the data obtained at the 256-cell stage. The expression levels of genes that are misregulated in *mir430*^-/-^ + inj MiR430 embryos at the 256-cell stage (with a median of 3.5 log_2_ fold difference with WT for the upregulated genes, and 2.9 log_2_ fold for the downregulated genes; Fig. 2d) gradually recover during consecutive developmental stages, reaching WT expression levels at Sphere stage (with a median of 0.03 log_2_ fold difference with WT for the upregulated genes, and 0.07 log_2_ fold for the downregulated genes; Fig. 2d). Looking at all genes, we find that overall fewer genes are misregulated at later stages, from 716 up and 242 down at 256-cell to 51 up and 30 down at Sphere stage (Extended Data Fig. 5). We conclude that in the absence of *mir430* transcription bodies, gene expression is misregulated, but that this effect is short-lived as expression levels return to WT levels in a timeframe of a couple of cell cycles. This likely explains the lack of obvious developmental defects in *mir430*^-/-^ + inj MiR430 embryos.

We then further analysed the genes that are misregulated at the 256-cell stage. The downregulation of genes suggests that transcription bodies facilitate transcription of genes other than *mir430* itself, consistent with work in which transcribed genes were shown to colocalize ^12–14,26^, as well as the proposed role of transcription bodies in increasing transcriptional efficiency ^18,27,28^. We hypothesize that the genes that are downregulated in *mir430*^-/-^ + inj MiR430 embryos localize to the *mir430* transcription bodies in WT embryos and are positively affected by this localization. The observation that downregulated genes return to WT expression levels over time suggests that transcription bodies that form during later stages in *mir430*^-/-^ + inj MiR430 embryos (Extended Data Fig. 6) may compensate for the loss of *mir430* transcription bodies. Here, we focus on the upregulated genes. Their high number suggest that the *mir430* transcription bodies sequester components of the transcriptional machinery, preventing genes elsewhere in the nucleus from being transcribed.

To investigate this sequestration model, we characterized the genes that are upregulated in response to *mir430* transcription body removal. We first asked when these genes are induced in WT embryos. We analysed their expression at the 1024-cell, Oblong, and Sphere stages, and compared this to their expression at the 256-cell stage. We found that most of the genes (81%) are activated during genome activation in WT embryos (Fig. 3a), indicating that loss of the *mir430* transcription bodies specifically activates the expression of genes that are about to be activated, with 62% (446/716) being advanced by 30 minutes (at the 1024-cell stage in WT, Fig. 3a). When focusing on the genes that are advanced by 30 minutes, we observe a good correlation between their degree of upregulation between *mir430*^-/-^ + inj MiR430 and WT at 256-cell stage, and between 1024-cell and 256-cell stage in WT (Fig. 3b and Extended Data Fig. 7). This further supports the notion that the effect on transcription activation is specific.

**Figure 3.**
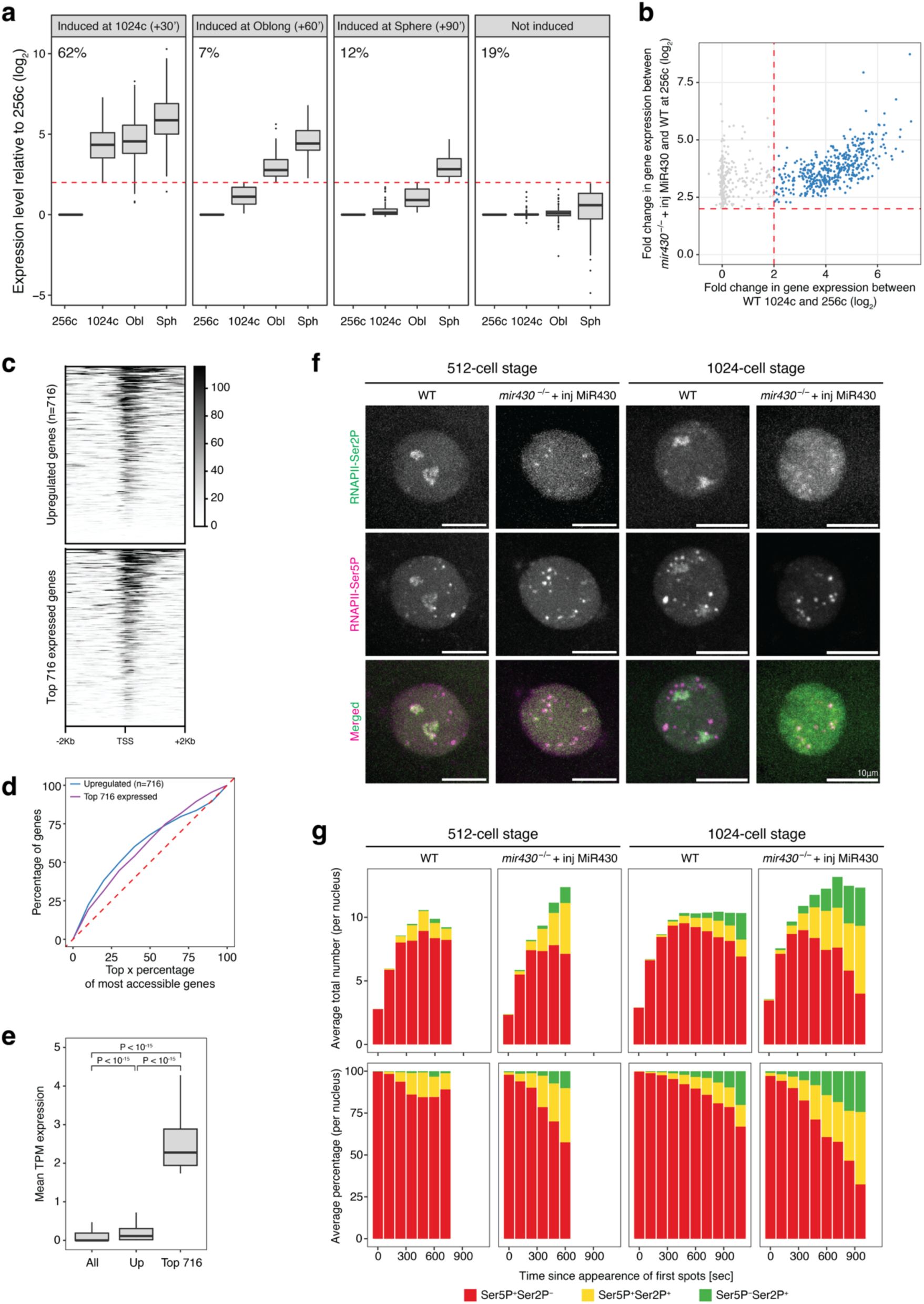
Loss of *mir430* transcription bodies causes premature gene activation. **a.** Time of induction in WT embryos for the 716 genes that are upregulated in *mir430*^-/-^ + inj MiR430 *vs.* WT embryos (at 256-cell stage). Expression at 256-cell is used as a reference. The genes are split in four groups based on when they are induced. The percentage of genes in each group is indicated. **b.** Scatterplot representing the fold change in expression between *mir430*^-/-^ + inj MiR430 and WT embryos at 256-cell on the y-axis, and between 1024-cell and 256-cell in WT embryos on the x-axis. All upregulated genes are shown in grey, genes induced at 1024-cell (62%) are shown in blue. **c.** Heatmap of chromatin accessibility at 256-cell of the 716 genes that are upregulated in *mir430*^-/-^ + inj MiR430 embryos compared to WT embryos at 256-cell stage, and – for comparison – the 716 most expressed genes in WT embryos at 256-cell stage. Genes are ranked by accessibility. **d.** Graph showing the fraction of promoters of the 716 upregulated (in *mir430*^-/-^ + inj MiR430 *vs.* WT at 256-cell stage), and the 716 most expressed genes (in WT at 256-cell) amongst the x% most accessible promoters at 256-cell in WT embryos. **e.** Average expression level of the 716 upregulated genes (*mir430*^-/-^ + inj MiR430 *vs.* WT at 256-cell stage), the 716 most expressed genes, and all annotated genes, at the 256-cell stage, in WT. Outliers are not shown. **f.** Visualization of RNAPII-Ser2P and RNAPII-Ser5P with antigen-binding fragments (Fabs) in WT and *mir430*^-/-^ + inj MiR430 embryos at 512- and 1024-cell stage. Shown are representative micrographs of individual nuclei. See Extended Data Fig. 9-10 for complete cell cycles. **g.** Quantification of Ser5P-positive/Ser2P-negative (red), Ser5P-positive/Ser2P-positive (yellow) and Ser5P-negative/Ser2P-positive (green) transcription bodies during the cell cycle at 512-cell and 1024-cell, as shown in f. Number (upper panel) and percentage (lower panel) are shown.

We then investigated the chromatin accessibility state of the genes that are upregulated. We analysed an ATAC-Seq dataset we had previously generated in WT embryos ^29^ and found that the transcription start sites of genes that are upregulated in the *mir430*^-/-^ + inj MiR430 are highly accessible in WT embryos (Fig. 3c), being as accessible as those of the most highly expressed genes (Fig. 3c-d), even though they are hardly expressed (Fig. 3e). This suggests that potentially sequestered factor(s) function downstream of transcription-factor binding and chromatin opening. Consistent with this, motif analysis of the promoters of upregulated genes did not identify transcription factor motifs that were both sufficiently enriched compared to unexpressed genes and present in a sufficiently high fraction of the upregulated genes to explain our observations (Extended Data Fig. 8). Even motifs of Nanog, Pou5f3 and Sox19b, transcription factors that have been shown to be required for the activation of transcription during ZGA in zebrafish ^9,29–32^ did not fulfil these two requirements. Taken together, our results suggest that the presence of the *mir430* transcription bodies prevents a significant number of genes that are poised for activation from going into elongation.

This reminded us of the observation that WT embryos from the 256-cell stage onward, in addition to the *mir430* transcription bodies in which both transcription initiation and elongation can be observed, display transcription bodies that are positive for transcription initiation, but not elongation ^9^. We were previously unable to explain this observation but now hypothesized that the *mir430* transcription bodies might sequester a pause release factor, causing genes in these transcription bodies to be arrested at the initiation phase. If this were true, these transcription bodies would be predicted to acquire elongation signals in nuclei of *mir430*^-/-^ + inj MiR430 embryos. To investigate this, we injected 1-cell WT and *mir430*^-/-^ + inj MiR430 embryos with fluorescently labelled antigen-binding fragments (Fabs) recognizing RNAPII-Ser5P (transcription initiation) and RNAPII-Ser2P (transcription elongation), as described before ^9,22,23^. Consistent with our hypothesis, we saw more Ser2P-positive transcription bodies in *mir430*^-/-^ + inj MiR430 than in WT embryos during the 512- and 1024-cell stages (Fig. 3f, and Extended Data Fig. 9-10). Quantification of initiation-only, initiation/elongation, and elongation-only transcription bodies confirmed that more transcription bodies go into productive elongation in *mir430*^-/-^ + inj MiR430 than in WT embryos (with a maximum of 15% elongation-positive transcription bodies in WT versus 33% in *mir430*^-/-^ + inj MiR430 at the 256-cell stage, and a comparable difference at the 1024-cell stage; Fig. 3g). Thus, the removal of the *mir430* transcription bodies causes transcription bodies that are arrested in the initiation phase to transition to the transcription elongation phase.

Our data suggests that transcription bodies sequester a pause release factor. Pause release is regulated by the pTEFb complex of which CDK9 is the catalytic subunit. We visualized CDK9 in WT embryos by immunofluorescence and found that it is strongly enriched in the two *mir430* transcription bodies (Fig 4a). If the amount of CDK9 in the nucleus were limiting and its sequestration in *mir430* transcription bodies would hamper transcription elongation elsewhere in the nucleus, it would be predicted that overexpression of CDK9 in WT embryos would induce the formation of ectopic transcription bodies. To directly test this, we compared the occurrence of ectopic transcription bodies between WT embryos with and without injected *cdk9* mRNA. We visualized RNAPII Ser2P to detect transcription bodies. To distinguish transcription bodies that are nucleated by the *mir430* locus from ectopic transcription bodies, we also visualized *miR430* transcripts using Morpholino Visualization of Expression (MOVIE ^21^). At the 128-cell stage, we detected two prominent *mir430* transcription bodies in nuclei of WT embryos, as well as in WT embryos that were injected with *cdk9* mRNA (Fig. 4b and Extended Data Fig. 11). In the latter, however, we detected additional transcription bodies. Because these lack *miR430* MOVIE signal and we do not detect them in uninjected WT embryos, we consider these to be ectopic transcription bodies. The ectopic bodies are small and limited in number at the 128- and 256-cell stage, and more prominent at the 512- and 1024-cell stage (Fig. 4b and Extended Data Fig. 11-14). The abundance of ectopic transcription bodies in WT embryos in which CDK9 is overexpressed, phenocopies the abundance of ectopic transcription bodies seen in *mir430*^-/-^ + inj MiR430 embryos. The quantification of ectopic transcription shows that at all four examined stages, ectopic transcription bodies are significantly more abundant in nuclei in which CDK9 is overexpressed (WT + inj *cdk9* mRNA), or in which the large *mir430* transcription bodies are disrupted (*mir430*^-/-^ + inj MiR430), than in WT embryos (Fig. 4c). This supports our hypothesis that CDK9 is sequestered in *mir430* transcription bodies, limiting CDK9 availability elsewhere in the nucleus.

Taken together, we show that the removal of the *mir430* transcription bodies results in a genome-wide misregulation of gene expression. Many genes are downregulated, consistent with a model where the bodies facilitate transcription by promoting the biomolecular interactions underlying gene expression ^3,16,18^. Unexpectedly, we also observed many genes that are upregulated upon disruption of the *mir430* transcription bodies. We show that transcription bodies sequester the pause release factor CDK9, preventing genes that are not present in the transcription bodies from being transcribed (Extended Data Fig. 15). Factor sequestration regulating biological processes has been described before ^33,34^ and a recent study showed that the loss of the transcription factor Zelda in *Drosophila* embryos caused RNAPII to relocate from Zelda-containing transcription bodies to the histone locus body ^35^. More generally, work in mouse embryos has shown that CDK9 abundance is an important factor in the timing of zygotic genome activation^36^. Our work shows that endogenous transcription bodies play an important role in controlling the availability of transcriptional machinery in the nucleus.

## Acknowledgements

We thank the members of the Vastenhouw lab for technical support, helpful feedback, and stimulating discussions. We thank the Pauli and Ameres labs for technical advice on SLAM-Seq. We thank Maciej Kerlin, Aleksandar Vještica, and Edlyn Wu for comments on the manuscript, and the following facilities and services for their support: MPI-CBG - fish facility, light microscopy facility, and scientific computing, UNIL - genomic technologies facility, cellular imaging facility and the bioinformatics competence center. M.U. was supported by the Boehringer Ingelheim Fonds. Research in N.L.V’s laboratory was supported by the Max Planck Society, the University of Lausanne, a European Research Council Consolidator Grant (101003023), the Volkswagen Foundation (94773), and the German Research Foundation (VA 1209/2-1). Research in H.K.’s laboratory was supported by the Japan Society for the Promotion of Science KAKENHI (JP18H05527, JP21H04764 and JP20K06484).

## Author contributions

Conceptualization: M.U., N.L.V. Methodology: M.U., K.K., H.U. Investigation: M.U., K.K. Data analysis: M.U. Writing – original draft: M.U. and N.L.V. Writing – reviewing & editing: all authors. Funding acquisition: M.U., N.L.V., H.K. Supervision: N.L.V.

## Data and materials availability

Raw sequencing data will be uploaded on Gene Expression Omnibus (GEO) prior to publication. Raw imaging data are available upon request. All other data is available in the main text or the extended data.

## Methods

### Zebrafish and molecular biology approaches

#### Zebrafish husbandry and manipulation

Zebrafish were maintained and raised under standard conditions, and according to Swiss regulations (canton Vaud, license number VD-H28). Wild type (TLAB) and mutant (*mir430*^-/-^) ^9^ embryos were collected immediately upon fertilization and allowed to develop to the desired stage at 28°C. *mir430*^-/-^ embryos were rescued by injecting 2nL of rescue solution (10μM of MiR430a, MiR430b and MiR430c duplex in 1x siRNA solution (300mM KCl, 30mM HEPES, 1mM MgCl_2_, brought to pH 7.5 with KOH)), as described before ^9,24^. If the chorion was chemically removed, this was done by incubating embryos right after fertilization in 1.5 mg/mL Pronase E (Sigma-Aldrich, #107433) for approximately five minutes. Developmental stage was determined by morphology (according to ^37^), the distance between nuclei, as well as cell size, cell cycle duration, and synchronicity ^9^. Fabs were injected into the cell at 0.44ng/embryo (αRNAPII-Ser2P-Alexa488), or 0.66ng/embryo (αRNAPII-Ser5P-Cy5). Lissamine-labelled anti-*miR430* morpholino (5’-TCTACCCCAACTTGATAGCACTTTC-3’-Lissamine, Gene Tools) was injected into the cell at 14 fmole/embryo ^21^. s^4^UTP (TriLink, tebu-bio GmbH, #N- 1025-1, diluted to 50mM in 10mM Tris-HCl pH 7.5) was injected into the yolk at 50nM/embryo.

#### Genotyping

Embryos (live and fixed) were genotyped using the HotSHOT method ^38^.

#### mRNA production and injection

eGFP with three perfect *miR430* target sites in its 3’UTR (eGFP-3xPT-miR430b) was inserted in the pCS2^+^ plasmid (Addgene) as described before ^24^. The coding sequence of *cdk9* (ENSDART00000065859.8) was amplified from cDNA using SuperScript III (Invitrogen, #18080085) and oligodT primers, and inserted in the pCS2^+^ plasmid (Addgene) using restriction-enzyme cloning. mRNA was synthesized using the mMessage mMACHINE Transcription Kit (ThermoFisher, #AM1340) and purified with the RNeasy MinElute Cleanup Kit (QIAGEN, #74204). mRNA was injected into embryos at the 1-cell stage in the following amounts: 50pg of eGFP mRNA, and 100pg of RFP or *cdk9* mRNA.

#### Preparation of antigen-binding fragments (Fabs)

Fluorescently labelled Fabs specific to RNAPII-Ser5P and RNAPII-Ser2P were prepared from monoclonal antibodies specific to RNAPII-Ser5P and Ser2 phosphorylation ^39–41^. Monoclonal antibodies were digested with Ficin (ThermoFisher Scientific), and Fabs were purified through protein A-Sepharose columns (Cytiva) to remove Fc and undigested IgG. After passing through desalting columns (PD MiniTrap G25; Cytiva) to substitute the buffer with PBS, Fabs were concentrated up to >1 mg/ml using 10 k cut-off filters (Amicon Ultra-0.5 10 k; Merck). Fabs were conjugated with Alexa Fluor 488 (Sulfodichlorophenol Ester; ThermoFisher Scientific) or Cy5 (N-hydroxysuccinimide ester monoreactive dye; Cytiv) to yield ∼1:1 dye:protein ratio. After the buffer substitution with PBS, the concentration was adjusted to ∼1 mg/ml.

#### Probe production

A DIG-labeled antisense probe against *goosecoid* (*gsc*) was generated by *in vitro* transcription from a linearized plasmid using 10x Transcription buffer (Roche, #11465384001), 10x DIG RNA labelling mix (Roche, #11277073910), T7 RNA polymerase (Roche, #10881775001) and RNase OUT (Invitrogen, #10777019). Probe was purified with the RNeasy MinElute Cleanup Kit (QIAGEN, #74204), diluted in Hybridization buffer and stored at -20°C.

#### Whole mount in situ hybridization

Whole mount *in situ* hybridization (WM-ISH) was performed as described before ^42^, with some differences. In brief, embryos were fixed at the desired stage in 4% (w/v) PFA in PBS pH 7.4 overnight at 4°C. Embryos were then manually dechorionated with sharp forceps (Dumont No. 5) in PBS with 0.1% Tween-20 (PBST) and dehydrated by rinsing them in 50% and then 100% methanol in PBST on a nutator. At this point, embryos could be stored for several weeks at - 20°C. To continue, embryos were gradually rehydrated on a nutator, incubated in Hybridization buffer (50% formamide, 1.3x SSC pH 5.0, 5mM EDTA, 50μg/mL torula yeast RNA, 0.2% Tween-20, 0.1% SDS, 100μg/mL Heparin) for 1 hour at 70°C, and incubated with anti-*goosecoid* DIG-labelled probe overnight at 70°C. The next day, the probe was washed off with Hybridization buffer and embryos were incubated for 15 minutes in 50% (v/v) Hybridization buffer in TBST (25mM Tris-HCl pH 7.8, 137mM NaCl, 2.7mM KCl, 0.1% Tween-20) at 70°C. After rinsing with TBST, embryos were incubated for 1 hour in TBST and then in Blocking Buffer (2% blocking reagent in maleic acid buffer pH 7.5 (Roche, #11096176001) and 10% sheep serum (Sigma Aldrich, #S2263) in TBST) at room temperature on a nutator. anti-DIG antibody (Roche, #11093274910) was added in a dilution of 1:2.000 and embryos incubated overnight at 4°C on a nutator. The next day, embryos were washed with TBST and then NTMT (100mM NaCl, 100mM Tris-HCl pH 9.5, 50mM MgCl2, 0.1% Tween-20), and stained in NTMT-NBT-BCIP solution (4.5μL/mL of NBT (Roche, #11383213001) and 3.5μL/mL of BCIP (Roche, #11383221001) in NTMT) in the dark. Once sufficiently stained, embryos were washed with PBST, and then in 100%, 50% and 30% methanol in PBST at room temperature on a nutator. Embryos were stored at 4°C in PBST for up to a few days before imaging.

#### Immunofluorescence

Dechorionated embryos were fixed at the desired stage in 4% (w/v) PFA in PBS pH 7.4 overnight at 4°C. Embryos were then dehydrated by incubating them in 30%, 50% and then 100% methanol in PBS and incubated overnight at -20°C. To continue, embryos were gradually rehydrated. All following steps were performed while rocking. Embryos were washed multiple times with PBST (0.8% Triton-X 100 in PBS) and incubated in blocking buffer (10% NGS (Sigma-Aldrich, #G6767) and 4% BSA (Sigma Aldrich, #A8022) in PBST) for four hours at room temperature. Primary antibodies were added in 1% NGS in PBST overnight at 4°C (anti-CDK9, clone C12F7, Cell Signaling Technology, monoclonal IgG antibody generated in rabbit, 1:200 dilution; anti-RNAPII-Ser2P, clone H5, ab24758, monoclonal IgM antibody generated in mouse, Abcam, 1:1000 dilution). The next day, embryos were washed with PBST and then incubated with secondary antibodies (anti-rabbit-IgG AlexaFluor488-conjugated, Thermo Scientific, #A-21206, 1:500 dilution; anti-mouse-IgM AlexaFluor594-conjugated, Thermo Scientific, #A-21044, 1:500 dilution) in 1% NGS in PBST overnight at 4°C. Embryos were then washed with PBST, manually deyolked using sharp forceps (Dumont No. 5), and mounted in Vectashield Antifade Mounting Medium with DAPI (Vector Laboratories, VC-H-1200-10) before imaging them on a Spinning Disk microscope.

### Enriched SLAM-Sequencing (eSLAM-Seq)

#### Rationale

Early embryos contain large amounts of maternally loaded RNA ^19^ which mask the nascent RNAs in total RNA sequencing approaches. We therefore developed a method to sensitively detect nascent transcripts. Methods to enrich nascent transcripts have previously been described. Typically, an uracil/UTP analog (EU/s^4^UTP) is injected at the 1-cell stage and is incorporated into newly made RNAs. Such labelled transcripts can then be biotinylated and enriched by pull-down ^20,43^. SLAM-Seq is another approach in which nascent transcripts are labelled by s^4^UTP incorporation, but here the label is used to convert 4-thio-uracils to carboxyamido-methylated 4-thio-uracils, which will be misread as a cytosine during reverse transcription, introducing specific point mutations that enable the identification of reads derived from labelled transcripts by sequencing ^44–46^. We combined these two approaches in enriched SLAM-Seq (eSLAM-Seq), which is conceptually similar to TT-TimeLapse-Seq and TT-SLAM-Seq ^47,48^. Specifically, we used a combination of s^4^UTP labelling ^43^ with improved biotinylation and enrichment chemistry ^49,50^, followed by SLAM-Seq chemistry.

#### Sample preparation

Because of the light-sensitive nature of s^4^UTP, all steps until the treatment of RNA with iodoacetamide (IAA) were performed in the dark. When staging embryos at the stereomicroscope, for example, we used the lowest light-setting that allowed us to still see the embryos, and a UV filter was set between the injection plate and the light source to protect the s^4^UTP. Successive steps were performed protecting the samples from the light by using low light settings in the laboratory room, by working inside a cardboard box, and by covering the samples with aluminium foil.

Synchronized s^4^UTP-injected embryos were collected and left to develop to the desired stage. They were collected in as little 0.3x Danieau’s Buffer as possible, homogenized in 500μL Qiazol (QIAGEN, #79306) with a 5mm stainless steel bead (QIAGEN, #69989) using a TissueLyser (30 seconds at 30Hz) and stored at -80°C. 100 embryos (256-cell stage) or 50 embryos (1024-cell, Oblong and Sphere stage) were used. Chloroform was added and RNA was extracted by phase separation, using a MaXtract tube (QIAGEN, #129046), after which it was precipitated with isopropanol supplemented with 0.2mM DTT and glycogen. The pellet was washed with 80% ethanol supplemented with 1mM DTT, air-dried, and resuspended in 40μL of 1mM DTT. DNase treatment was performed in 100μL of 1x DNAse reaction buffer with 4U of TURBO DNase (ThermoFisher, AM2238) for 30 minutes at 37°C. The volume was brought to 250μL with nuclease-free water and RNA was extracted with 25:24:1 Phenol-Chloroform-Isoamyl alcohol (Invitrogen, #15593-031) using a MaXtract tube, precipitated with isopropanol supplemented with 0.2mM DTT, glycogen and sodium acetate. The pellet was washed with 80% ethanol supplemented with 1mM DTT, air-dried, and resuspended in 40μL of nuclease-free water.

Labelled RNA was biotinylated in 100μL of 20mM HEPES pH 7.4, 1mM EDTA pH 8.0, 10ng/μL MTSEA-biotin-XX (Biotium, #90066) and 20% DMF) by incubation at room temperature for 30 minutes while rotating. The volume was brought to 250μL with nuclease-free water and RNA was extracted with 24:1 Chloroform-Isoamyl alcohol (Sigma-Aldrich, #227056) using a MaXtract tube, and precipitated with isopropanol supplemented with glycogen and NaCl. The pellet was washed with 80% ethanol, air-dried, and resuspended in 50μL of nuclease-free water.

To enrich for biotinylated RNA, Dynabeads MyOne Streptavidin C1 beads (Invitrogen, #65001) were used. Per sample, 10μL of beads were washed twice in nuclease-free water and twice in high-salt wash buffer (100mM Tris-HCl pH 7.4, 10mM EDTA pH 8.0, 1M NaCl, 0.05% Tween-20), incubated for at least 1 hour at room temperature in bead-blocking buffer (40ng/μL glycogen in high-salt wash buffer) and washed twice with high-salt wash buffer. RNA was denatured at 65°C for 10 minutes, incubated for 5 minutes on ice and then added to the blocked beads. After incubation for 15 minutes while rotating at 30 rpm, the supernatant was removed and beads were washed 3 times with high-salt wash buffer, twice with 5.9M guanidinium chloride (with 5-minute incubation at room temperature between consecutive washes), once with TE buffer and three times with TE buffer (with 5-minute incubation at 55°C between consecutive washes). Bead-bound RNA was finally released by adding 25μL of freshly made elution buffer (100mM DTT, 20mM HEPES pH 7.4, 1mM EDTA pH 8.0, 100mM NaCl, 0.05% Tween-20) twice. Volume was brought to 100μL with nuclease-free water and RNA was precipitated with isopropanol supplemented with 0.2mM DTT, glycogen and NaCl. The pellet was washed with 80% ethanol supplemented with 1mM DTT, air-dried, and resuspended in 20μL of 1mM DTT. Iodoacetamide-mediated alkylation was performed in 10mM IAA (freshly dissolved in 100% ethanol, BioUltra, #I1149), 50mM NaPO_4_ pH 8.0 and 50% DMSO for 15 minutes at 50°C. Samples were then transferred on ice and the reaction stopped by quickly adding 1.4μL of 1M DTT. RNA was precipitated with 100% ethanol supplemented with glycogen and sodium acetate overnight. The pellet was washed with 80% ethanol, air-dried, and resuspended in 8.5μL of nuclease-free water.

Libraries were prepared with the TruSeq stranded mRNA Kit (Illumina, #20020594) and TruSeq RNA UD Indexes (Illumina, #20022371). The polyA enrichment step was not performed. Fragmentation was performed for 1 minute at 94°C. Paired end 150bp sequencing was performed on an Illumina NovaSeq 6000 System, loading each library on two lanes (Illumina Reagent Kit v1.5). Sequencing data were demultiplexed with bcl2fastq 2.20.

#### Quality controls

To assess the integrity of total RNA (Extended Data Fig. 4c, left panel), 1μg/sample was diluted in 2x RNA Gel Loading Dye (Thermo Scientific, #R0641), denatured for 90 seconds at 85°C, chilled for 5 minutes on ice and loaded onto an agarose gel alongside RiboRuler High Range RNA Ladder (ThermoFisher, #SM1823). The presence of two prominent bands (rRNA) are an indicator of RNA integrity.

To test the specificity of s^4^UTP biotinylation (Extended Data Fig. 4c, right panel), the RNA was then transferred from the gel onto a Nylon Hybond-N+ membrane (Amersham, #RPN203B) by capillary transfer using a Whatman Turboblotter in 20X SSC according to manufacturer’s instructions overnight. After drying, the membrane was crosslinked twice with 120mJ/cm^2^ using a UV Stratalinker 1800 (Stratagene). The membrane was then incubated for 20min in Blocking Buffer (10% SDS and 1mM EDTA in 1xPBS pH 7.5), for 1h in 1:10.000 Streptavidin-HRP conjugate (Perkin, #NEL750) in Blocking buffer, 2x for 10min in 10% SDS in 1xPBS pH 7.5, 2x for 10min in 1% SDS in 1xPBS pH 7.5, and 2x for 10min in 0.1% SDS in 1xPBS pH 7.5, all while nutating ^51^. Finally, the signal was visualized with ECL Western Blotting Detection Reagents (Amersham, #RPN2106) according to manufacturer’s instructions, and documented with a Fusion FX (Vilber).

### Bioinformatic analyses

#### eSLAM-Seq

To map the raw sequencing data we concatenated Fastq files that came from the same library but that were run on different lanes with the ‘cat’ command (GNU core utilities, linux terminal), trimmed them using Trimmomatic (Paired-end mode, LEADING:3 TRAILING:3 SLIDINGWINDOW:4:15 MINLEN:36) ^52^, and mapped them using HISAT-3N (--base-change T,C --repeat --rna-strandness RF --fr --no-discordant) ^53^, which is a HISAT variant specifically designed for methods like SLAM-Seq that introduce specific point-mutations. Uniquely mapped fragments were filtered with Samtools ^54^ and mate information was fixed with Picard FixMateInformation (--IGNORE_MISSING_MATES FALSE) ^55^.

To distinguish between naturally occurring SNPs and SLAM-Seq-specific mismatches, we first identified naturally occurring SNPs by analysing four IAA-minus samples (one per stage) in which the iodoacetamide treatment – and therefore the base conversion – was not performed. The bam files were converted to a suitable format with the Samtools mpileup function and then SNPs were identified by Varscan pileup2snp (--min-coverage 20 --min-reads2 5 --min-avg-qual 15 --min-var-freq 0.25 --p-value 0.01). The union of the SNPs that were identified was used as a consensus list. Then the splbam.py script from pulseR ^56^ was used to select fragments with at least one SLAM-Seq-specific mismatch.

To assess gene expression levels of annotated genes, raw read counts were obtained using FeatureCounts (-p --countReadPairs -B --minOverlap 10 -s 2 -t exon -g gene_id) ^57^. Normalization, Principal Component Analysis (PCA) and call for differential expression (alpha = 0.01 and log_2_(Fold Change) threshold = 2) was performed using DESeq2 ^58^. When performing PCA, only the top 500 genes (selected by highest variance in expression levels) were considered. When counting the number of differentially expressed genes, mitochondrial and *mir430*-coding genes were excluded.

#### Motif analysis

To assess the presence and enrichment of specific motifs in promoters of upregulated genes, we started with the non-redundant JASPAR 2022 Vertebrate core database. From this, only motifs whose cognate transcription factors are expressed during ZGA (Riboprofiling dataset, >10rpkm, ^30^) were kept. The Nanog motif ^29^ and Pou5f1 motif ^31^ were added, because it is known that these factors are important during zebrafish ZGA. Next, we extracted promoter sequences (4000bp centred around the TSS) for upregulated genes as well as non-expressed genes (less than 10 raw reads in total across all samples, background group). Motif enrichment was then assessed using SEA (MEME Suite, ^59^). To estimate depletion of motifs, the background and input sample were inverted. Only motifs with a P-value < 0.05 and an E-value ≤ 10 were considered.

#### Promoter accessibility analysis

To analyse promoter accessibility of the upregulated genes in WT, we used an ATAC-Seq dataset that we previously generated ^29^. Replicates were merged and the signal in 4000bp centred around the TSS was plotted using Deep Tools ^60^. Promoter signal was used to rank all promoters by accessibility and to assess their enrichment amongst the most accessible promoters. The 716 most expressed (mean TPM) genes (excluding mitochondrial genes and *mir430*-coding genes) were used as a comparison.

#### Graph plotting for eSLAM-Seq

Plots were generated using R version 4.0.5 ^61^ with the ggplot2 ^62^ and Ggbio package ^63^.

### Microscopy

#### Fluorescence microscopy

To determine MiR430-activity using the GFP sensor assay, injected embryos were manually dechorionated at 24hpf and kept in Petri dishes. Fluorescence signal was measured on an OLYMPUS SZX16 stereo microscope with a MicroPublisher 5.0 RTV-R-CLR-10C colour camera (5x magnification, 400ms exposure time).

#### Bright field imaging

For bright field time lapse imaging, a mold ^64^ was used to create agarose wells using as little as possible 1% agarose in 0.3x Danieau’s solution in an Ibidi μ-Dish 35mm, high (#81158). The dish was mounted on a 28°C heated stage (by a Warner automatic temperature controller) and dechorionated embryos were loaded to the single wells at the 2-cell stage. Movies with 10-minute time intervals were taken on a Zeiss Axio Observer.Z1 microscope (Zeiss 5x/0.25 Fluar air objective, Zeiss AxioCam MRm Monochrome CCD camera). A z-stack (400μm in 9 steps) was taken for each embryo. A lid was used to reduce evaporation of buffer during imaging.

To assess developmental progression of live embryos past somitogenesis, snapshots of manually dechorionated embryos were taken at 24hpf, 48hpf and 72hpf on a Zeiss Axio Observer.Z1 microscope (Zeiss 2.5x/0.08 EC Plan-Neofluar air objective, Zeiss AxioCam ICc 1).

The gene expression pattern in fixed embryos after WM-ISH was visualized on a LEICA M165C stereomicroscope with LEICA MC170 HD colour camera (4x magnification). Embryos were oriented such that the dorsal region of the embryo was visible.

#### Spinning Disk microscopy

For imaging on a spinning disk microscope, dechorionated embryos were mounted at 32-cell stage in small droplets of 1% low-melting agarose (Invitrogen, #16520-050) in 75% v/v 0.3x Danieau’s solution and 25% v/v Optiprep Density Gradient Medium (Sigma Aldrich, #D1556)^65^ on a Ibidi μ-Dish 35mm, high (#81158). Embryos were brought closer to the coverslip surface by keeping the dish upside down until the agarose solidified (∼20 minutes). More agarose was then added to avoid drying out.

Transcription bodies in live embryos were imaged on a Nikon eclipse Ti2 with a Yokogava CSU-W1 Spinning Disk unit at 28°C (Okolab temperature control and microscope enclosure). Multi-position, z-stack (20μm in 41 steps, NIDAQ Piezo Z) time-lapse (2 minutes time interval) imaging was performed with a Nikon 60x/1.2 Plan Apochromat VC water objective. Images were captured in parallel on two Photometrics Prime 95B cameras with 100ms exposure time at 10% laser intensity.

Immunofluorescence signal (CDK9, RNAPII-Ser2P and DAPI) was also imaged on a Nikon eclipse Ti2 with a Yokogava CSU-W1 Spinning Disk unit. Images were taken with a Nikon 60x/1.2 Plan Apochromat VC water objective. Each channel was captured in series on a Photometrics Prime 95B camera with 100ms exposure time at 10% laser intensity each.

### Image processing and analysis

#### Software used for image analysis

Microscopy image handling was done using FIJI ^66^. Further data processing was carried out using R version 4.0.5 ^61^ and plots were generated with the ggplot2 package ^62^.

#### Analysis of eGFP reporter and RFP control

To assess *mir430* activity, mean fluorescence of eGFP (targeted by MiR430) and RFP (not targeted, and used as a normalizer) were measured in a rectangle of equal size in the head region and a background region outside of the embryo. The signal ratio was then calculated as follows: (eGFP_head_ – eGFP_background_) / (RFP_head_ – RFP_background_).

#### Detection and classification of mir430 and ectopic transcription bodies

We identified transcription bodies by the presence of RNAPII-Ser2P signal. To classify them into *mir430* and ectopic transcription bodies, we analysed *miR430* MOVIE signal. In case of overlap between RNAPII-Ser2P and MOVIE signal, bodies were classified as *mir430* transcription bodies. In case of no overlap between RNAPII-Ser2P and MOVIE signal, bodies were classified as ectopic transcription bodies (Extended Data Fig. 16). We used FIJI plugin ComDet ^67^, which detects spots using consistent intensity thresholding parameters that were manually set and used for all experiments. We note that frames taken after the disassembly of the nuclear envelope were not analysed because at that time transcription bodies also disassembled.

#### Detection and classification of transcription bodies by transcription state

Nuclei were segmented using the RNAPII-Ser5P channel, and RNAPII-Ser5P and Ser2P bodies detected using ComDet. This was used to classify transcription bodies into initiating, initiating-elongating and elongating bodies. We then identified and removed *mir430* transcription bodies from the analysis based on their morphological features (early appearance during cell cycle, larger size, growth behaviour) using the RNAPII-Ser2P channel. Here too, frames taken after the disassembly of the nuclear envelope were not analysed. For each nucleus, t=0 is set as the first frame at which spots appear, and the last time point is set as the last frame before prophase. Nuclei in which spots were detected for only one frame were discarded.

### Sample size and statistics

A minimum of 3 biological replicates (N) was acquired for each experimental condition. Lower-case n (n) refers to the number of embryos or nuclei for each experimental condition. Kruskal-Wallis tests were performed to determine if there is a statistically significant difference between the three genotypes in the degree of MiR430 activity as determined by the eGFP-Sensor assay (Fig. 1e), the time of completion of Epiboly (Fig. 1g) and the time of appearance of Kupffer’s vesicle (Fig. 1h). If this test was statistically significant (P-value < 0.05), pairwise comparisons with Bonferroni correction were performed using a pairwise Wilcoxon rank-sum test. A comparison was considered significant when adjusted P-value < 0.05. Dotplots were used to display the data, and adjusted P-values were reported using WT as reference. Red rhombuses represent the median.

A pairwise Wilcoxon rank-sum test with Bonferroni correction was performed to determine if the upregulated genes and the top 716 most expressed genes are significantly higher expressed compared to all genes (based on TPM expression values) (Fig. 3e). A comparison was considered significant when adjusted P-value < 0.05.

A pairwise Wilcoxon rank-sum test with Bonferroni correction was performed at each stage to determine if there is a statistically significant difference in the number of detected ectopic transcription bodies between WT, *mir430*^-/-^ + inj MiR430 and WT + inj *cdk9* mRNA (Fig. 4c). A comparison was considered significant when adjusted P-value < 0.05.

---

**Extended Data Figure 1.**
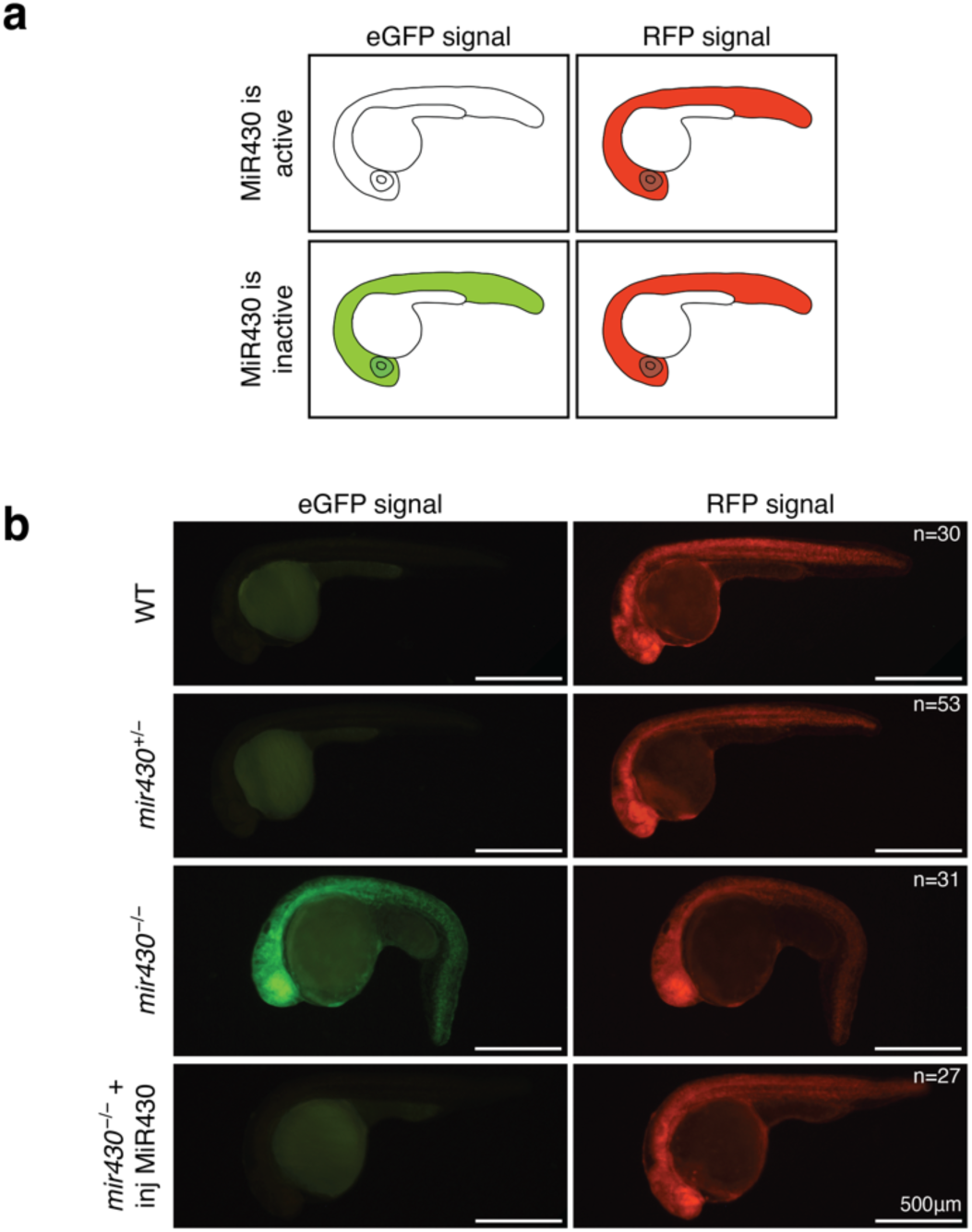
MiR430 activity is restored by injecting MiR430 RNA in *mir430*^-/-^ embryos. **a.** Schematic representation of expected eGFP and RFP expression in embryos with active and inactive MiR430 microRNA activity. **b.** Representative micrographs showing eGFP and RFP expression in WT, *mir430*^+/-^, *mir430*^-/-^ and *mir430*^-/-^ + inj MiR430 embryos at 24hpf.

**Extended Data Figure 2.**
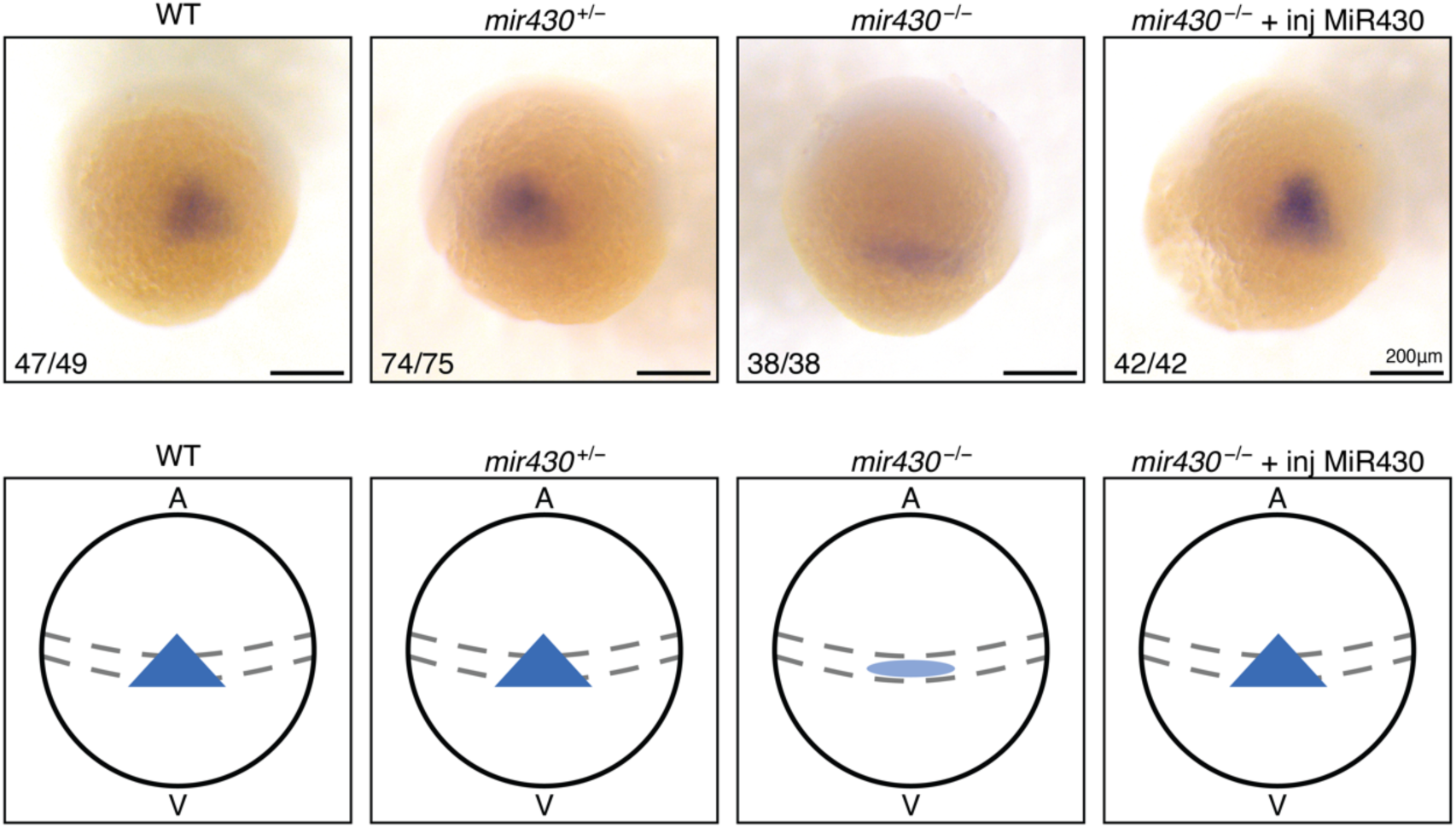
Phenotypic analysis of *mir430*^-/-^ embryos with and without injected MiR430 (*gsc* expression). Whole mount *in situ* hybridization (WM-ISH) of *Goosecoid (Gsc)* in WT, *mir430*^+/-^, *mir430*^-/-^ and *mir430*^-/-^ + inj MiR430 embryos at Shield stage. Representative micrographs show the embryos in dorsal view. Misexpression (reduced expression levels and reduced anterior extension of expression pattern) of *Gsc* can be observed in *mir430*^-/-^ embryos. The fraction in the lower left corner represents the frequency with which the shown phenotype was observed. A schematic representation of the phenotypes is shown below.

**Extended Data Figure 3.**
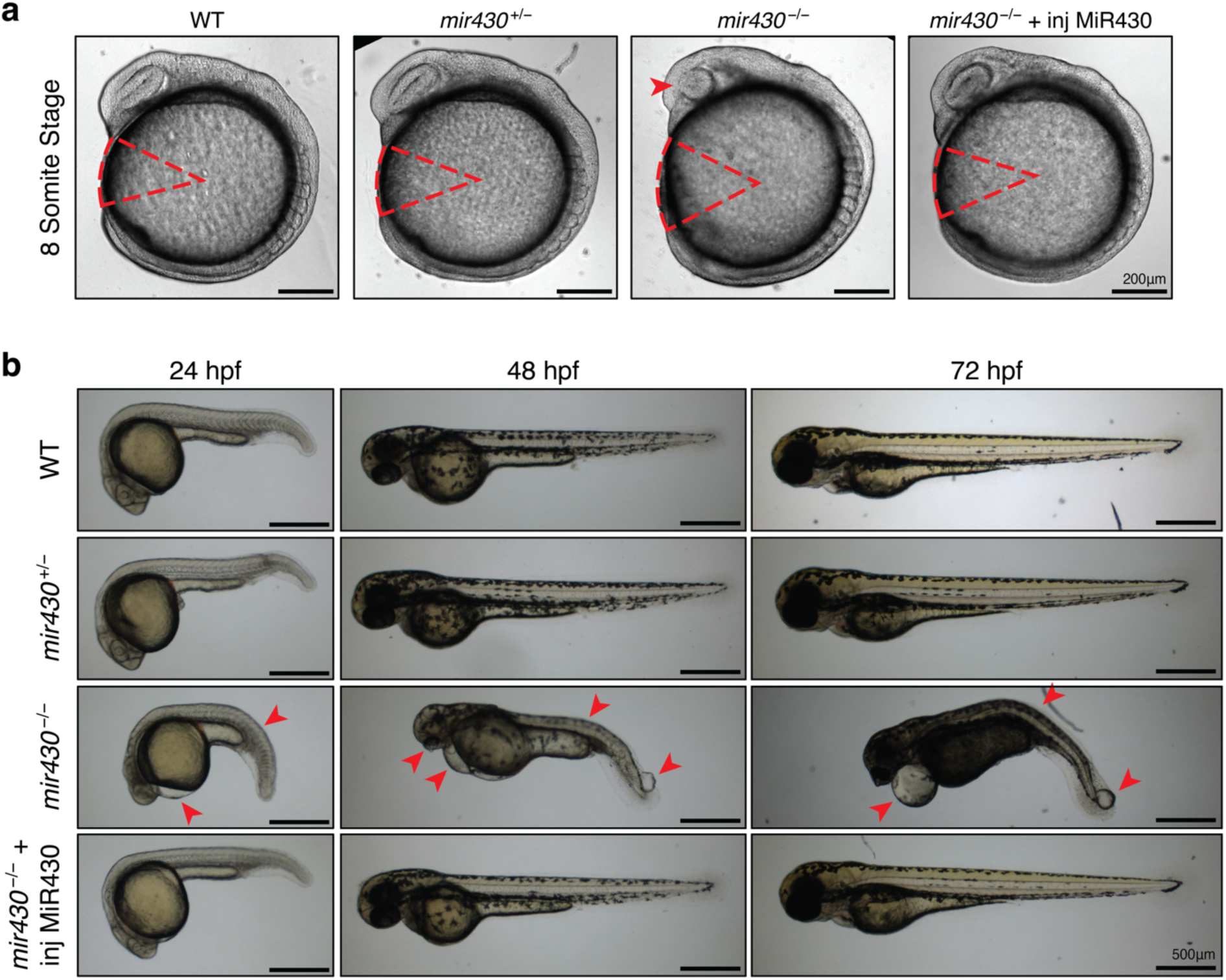
Phenotypic analysis of *mir430*^-/-^ embryos with and without injected MiR430 (developmental progress). **a.** Embryos at 8-Somite stage for WT, *mir430*^+/-^, *mir430*^-/-^ and *mir430*^-/-^ + inj MiR430 embryos. The reduced body axis extension and the malformation of the optic primordium in *mir430*^-/-^ embryos are highlighted. Representative micrographs are shown. **b.** Larvae at 24, 48 and 72 hours post fertilization (hpf) for WT, *mir430*^+/-^, *mir430*^-/-^ and *mir430*^-/-^ + inj MiR430 embryos. The malformation of trunk morphology and of the eye, the development of heart oedema and the appearance of blisters at the tail tip in *mir430*^-/-^ embryos are highlighted. Representative micrographs are shown. The 48hpf stage is also shown in Fig. 1i.

**Extended Data Figure 4.**
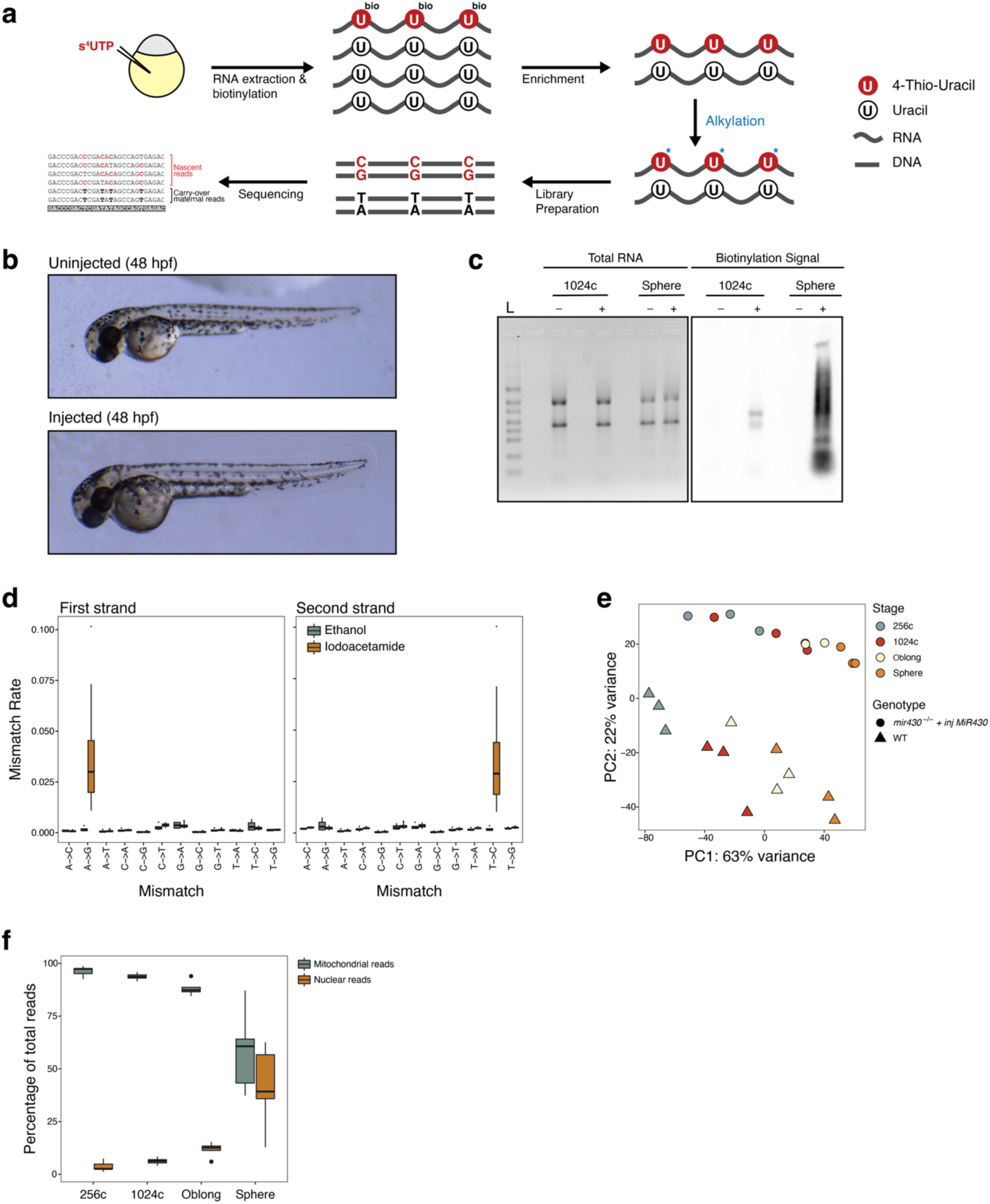
Establishment of enriched SLAM-Sequencing (eSLAM-Seq) in zebrafish embryos. **a.** Schematic representation of eSLAM-Seq protocol. See Methods for more detail. **b.** Representative micrographs of larvae that were/were not injected with 50nM/embryo of s^4^UTP at the 1-cell stage. No developmental malformations were observed upon injection. **c.** Equal amounts of biotinylated and heat-denatured total RNA derived from injected (+) and uninjected (-) embryos at 1024-cell and Sphere stage were run on an agarose gel (left panel). The RNA was then transferred on a nylon membrane and biotinylation detected by Streptavidin-HRP (right panel). Biotinylation was only observed in RNA derived from injected embryos, showing that biotinylation is specific for s^4^UTP-labeled RNA. L: RNA Ladder. **d.** Box plot representation of mismatch rate in Iodoacetamide- (N=24) and Ethanol-treated (N=4) samples. As expected, only Iodoacetamide-treated samples show a high level in A>G (First strand) and T>C (Second strand) mismatch rate. **e.** Principal Component Analysis (PCA) plot based on the top 500 genes (selected by highest variance in expression levels). Samples separate according to their developmental stage along the first PC and according to their experimental condition (WT and *mir430*^-/-^ + inj MiR430) along the second PC. **f.** Box plot representation of percentage of mitochondrial and nuclear reads, separated by developmental stage. Most reads are of mitochondrial origin at early ZGA, as shown before ^43^, with a gradual increase in the nuclear fraction during later stages.

**Extended Data Figure 5.**
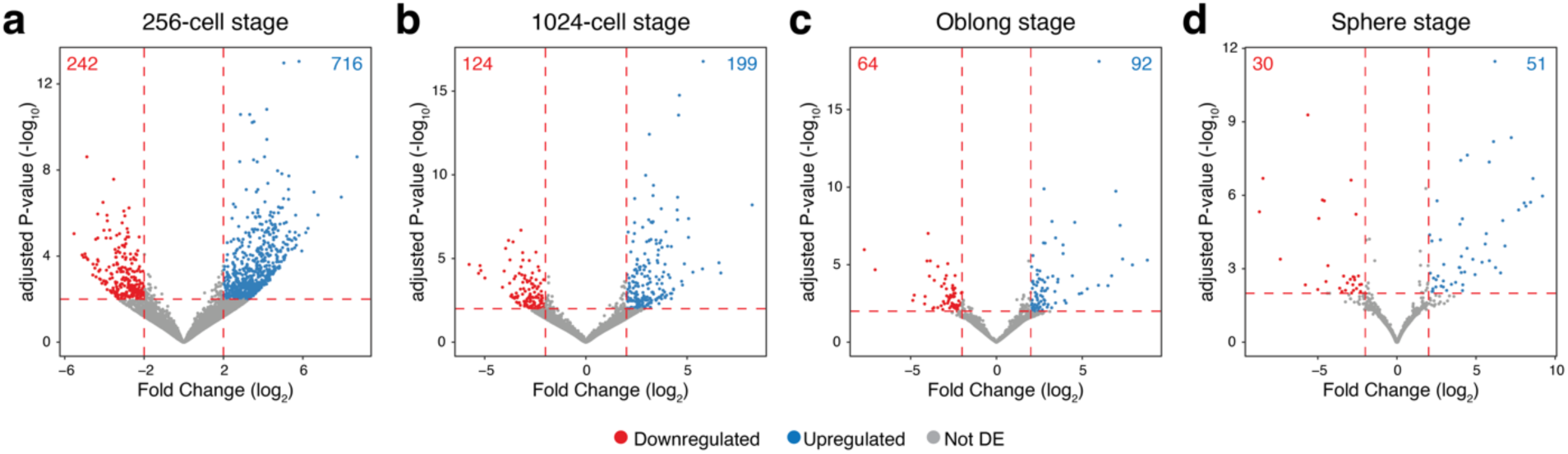
The disruption of two large endogenous transcription bodies causes short-term misregulation of gene expression. Volcano plots showing up- and downregulated genes in *mir430*^-/-^ + inj MiR430 *vs.* WT embryos at 256-cell **(a)**, 1024-cell **(b)**, Oblong **(c)** and Sphere **(d)** stage. Panel A is also shown in Fig. 2a.

**Extended Data Figure 6.**
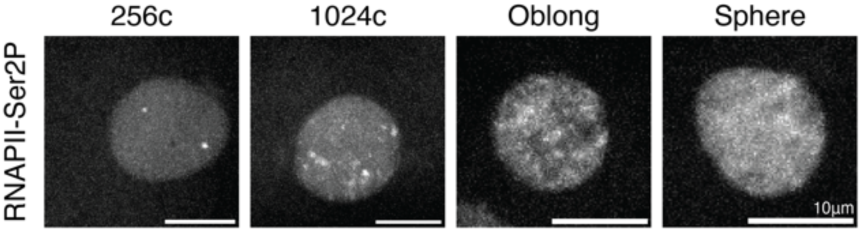
RNAPII-Ser2P signal at 256-cell, 1024-cell, Oblong and Sphere stage in *mir430*^-/-^ + inj MiR430 embryos. Visualization of RNAPII-Ser2P signal with Fabs. Representative micrographs are shown.

**Extended Data Figure 7.**
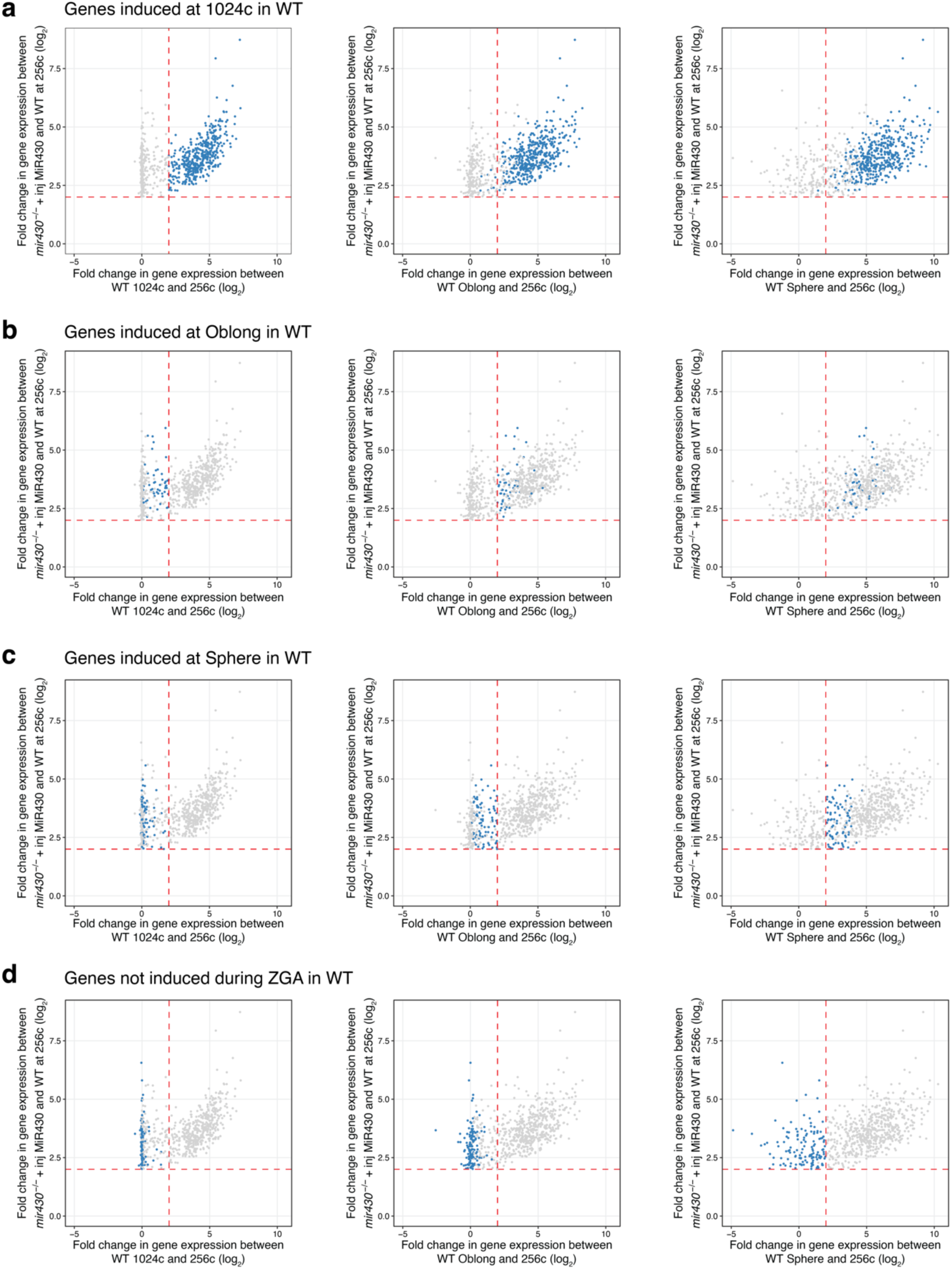
Correlation between expression level in *mir430*^-/-^ + inj MiR430 *vs.* WT and 1024-cell, Oblong and Sphere *vs.* 256-cell in WT embryos. **a.** Scatterplots representing the fold change in gene expression (log_2_ scale) between *mir430*^-/-^ + inj MiR430 and WT at 256-cell (y-axis), and the fold change in gene expression (log_2_ scale) between 1024- cell and 256-cell (left), Oblong and 256-cell (central) and Sphere and 256-cell (right) in WT embryos (x-axis). All upregulated genes are shown in grey, and genes induced at 1024-cell are highlighted in blue. The left graph is also shown in Fig. 3b. **b-d.** Same as in (A) but genes that are highlighted represent genes that are induced at Oblong **(b)**, Sphere **(c)**, or not induced during ZGA **(d)**.

**Extended Data Figure 8.**
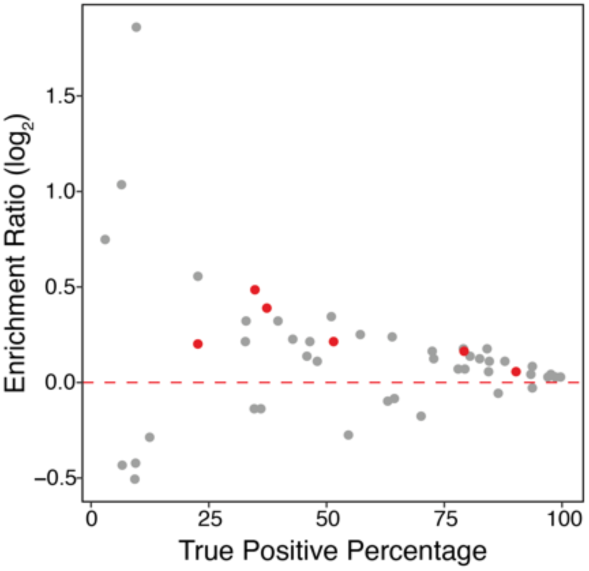
Transcription factor motif analysis of upregulated genes. Scatter plot representing the Enrichment Ratio (log_2_) of transcription factor motifs in the promoters (TSS ± 2Kb) of upregulated genes compared to the promoters of not expressed genes (y-axis), and the percentage of promoters of the upregulated genes that have the motif (x-axis). Only motifs whose corresponding protein is translated during early development ^30^ were considered, and only motifs with a p-value < 0.05 and E-value ≤ 10 are shown. Motifs corresponding to the three pluripotency factors Nanog, Sox19b and Pou5f3 (POU5F1, Pou5f1, Pou5f1::Sox2, POU5F1B, POU2F1, POU2F1::SOX2) are labelled in red.

**Extended Data Figure 9.**
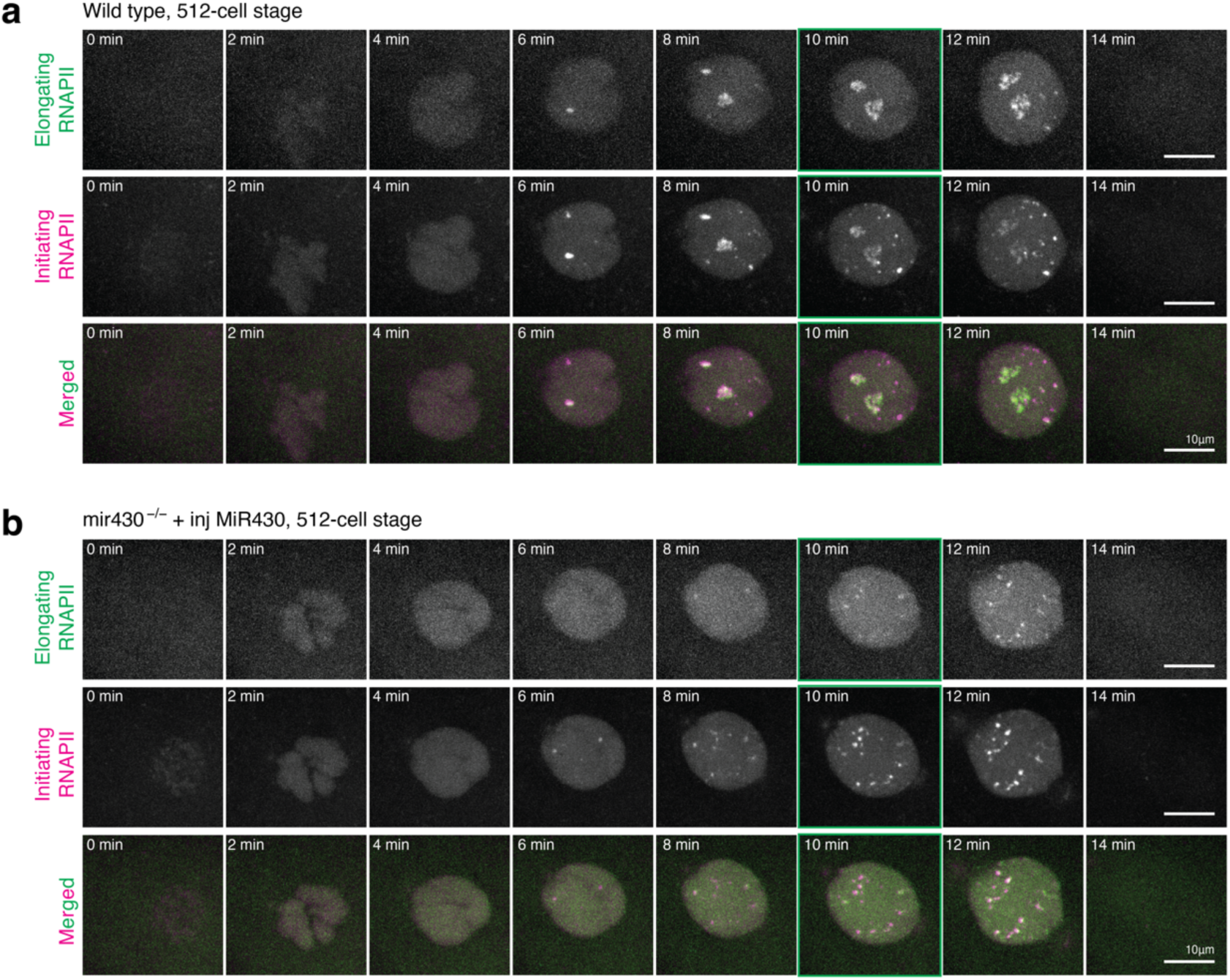
Stalled transcription bodies go into productive transcription in *mir430*^-/-^ + inj MiR430 embryos (512-cell stage). Visualization of initiating (Ser5P) and elongating (Ser2P) RNAPII with Fabs in WT **(a)** and *mir430*^-/-^ + inj MiR430 **(b)** embryos at 512-cell stage. Shown are representative images of individual nuclei, extracted from a spinning disk confocal microscopy time lapse. The micrographs shown in the main figure are highlighted in green.

**Extended Data Figure 10.**
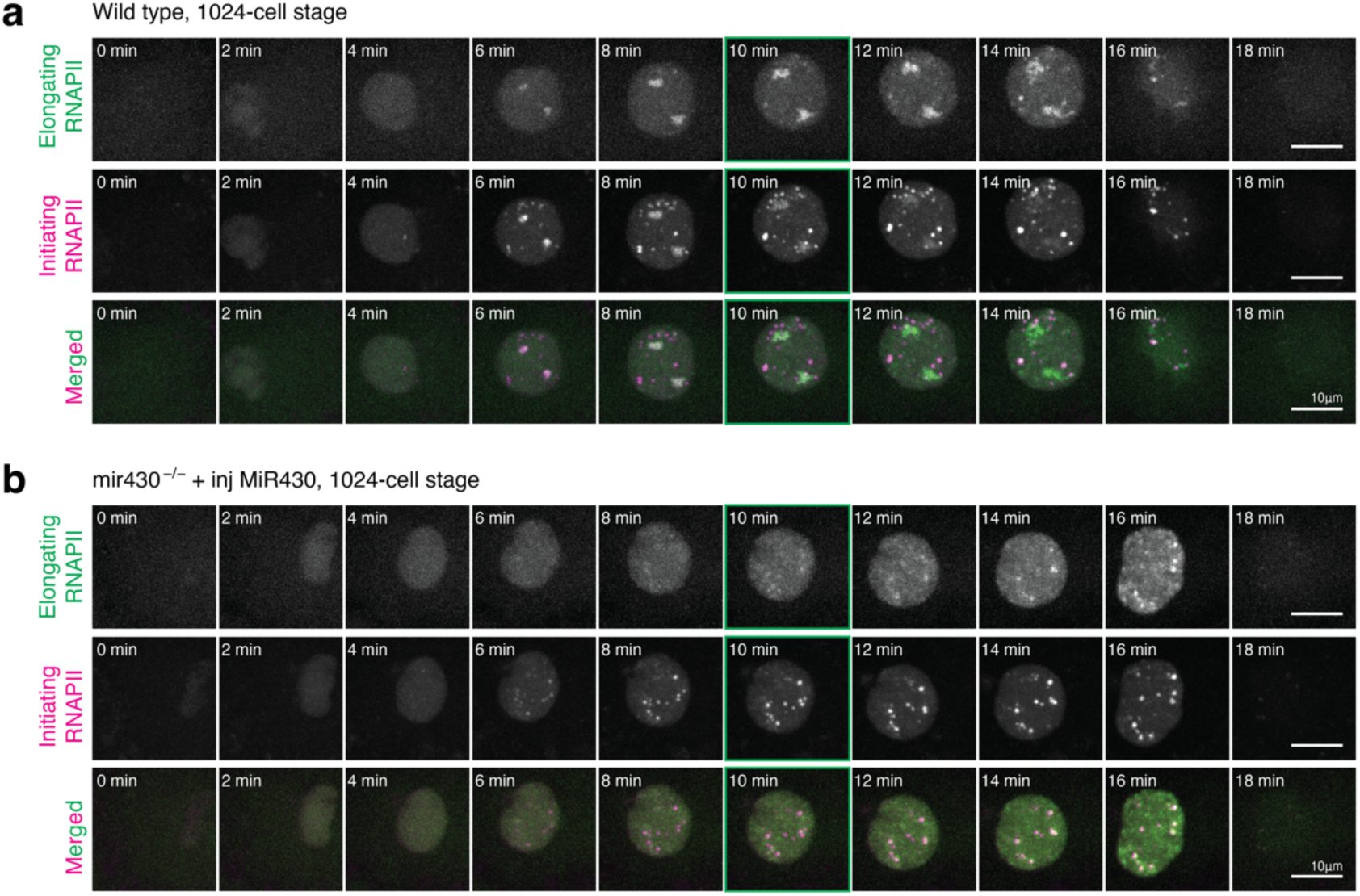
Stalled transcription bodies go into productive transcription in *mir430*^-/-^ + inj MiR430 embryos (1024-cell stage). Visualization of initiating (Ser5P) and elongating (Ser2P) RNAPII with Fabs in WT **(a)** and *mir430*^-/-^ + inj MiR430 **(b)** embryos at 1024-cell stage. Shown are representative images of individual nuclei, extracted from a spinning disk confocal microscopy time lapse. The micrographs shown in the main figure are highlighted in green.

**Extended Data Figure 11.**
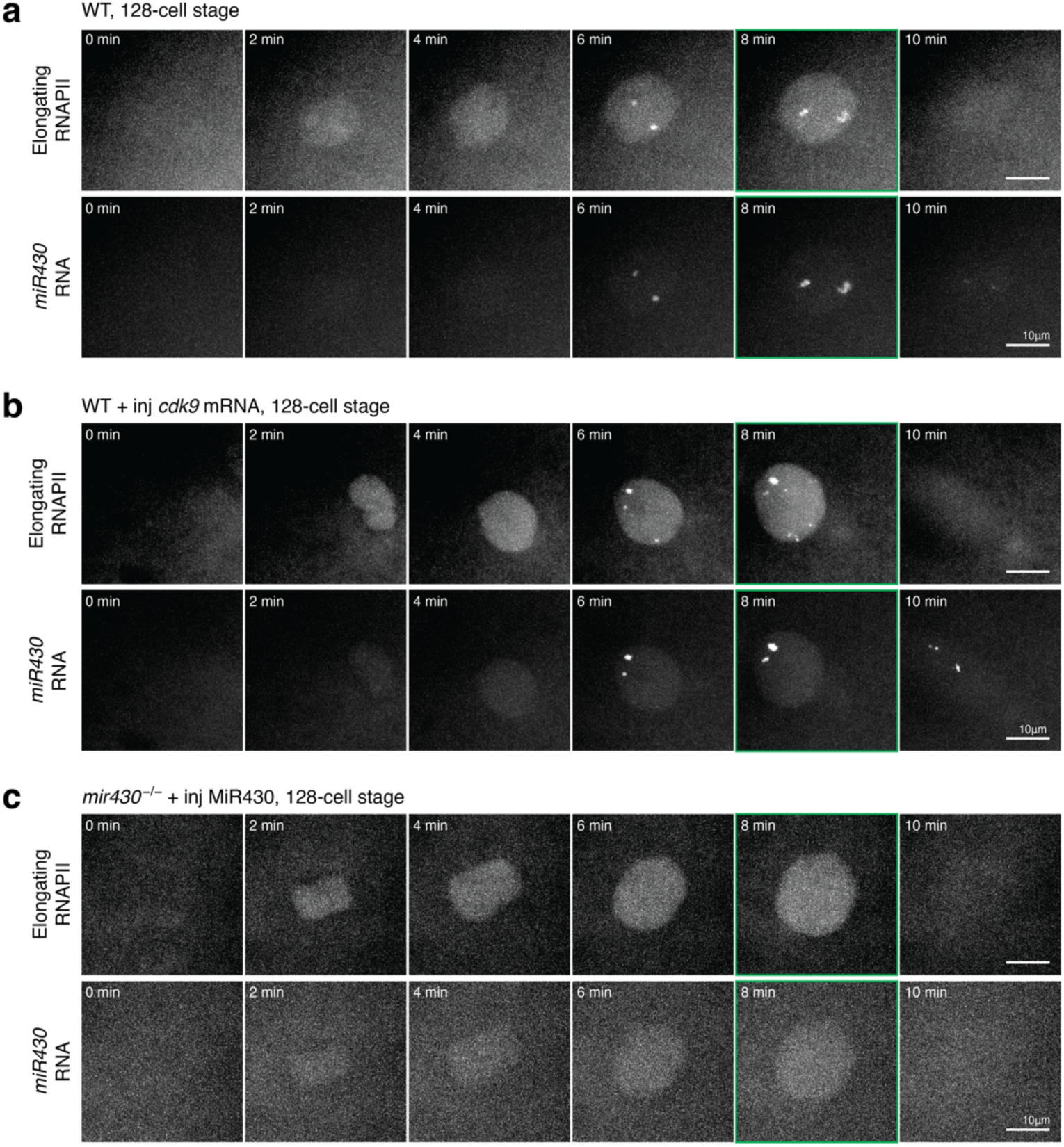
RNAPII-Ser2P signal in WT, WT + inj *cdk9* mRNA and *mir430*^-/-^ + inj MiR430 embryos at 128-cell stage. Visualization of RNAPII Ser2P (with Fabs) and *miR430* RNA (with MOVIE) in WT **(a)**, WT + inj *cdk9* mRNA **(b)**, and *mir430*^-/-^ + inj MiR430 **(c)** embryos at 128-cell stage. Shown are representative images of individual nuclei, extracted from a spinning disk confocal microscopy time lapse. The micrographs shown in the main figure are highlighted in green.

**Extended Data Figure 12.**
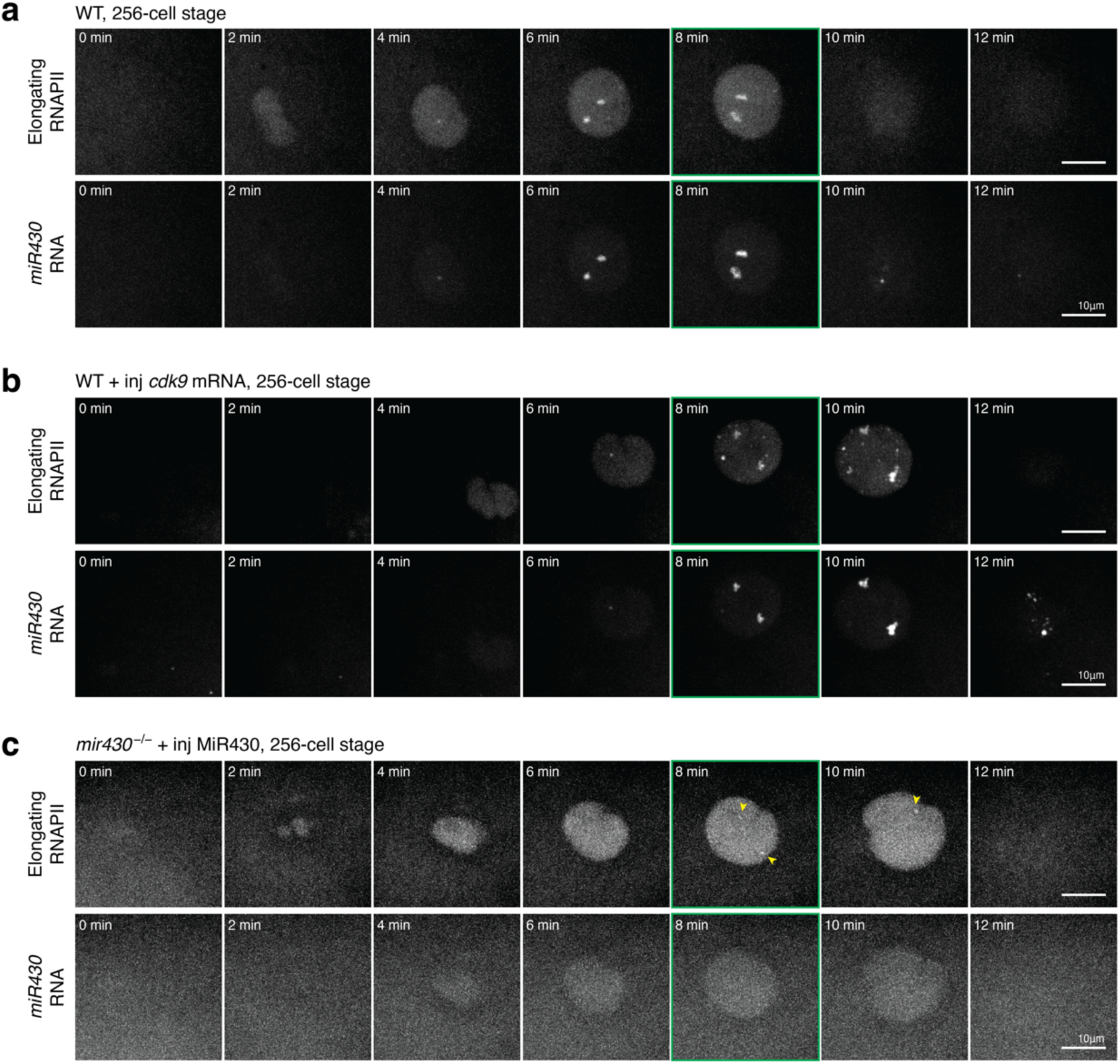
RNAPII-Ser2P signal in WT, WT + inj *cdk9* mRNA and *mir430*^-/-^ + inj MiR430 embryos at 256-cell stage. Visualization of RNAPII Ser2P (with Fabs) and *miR430* RNA (with MOVIE) in WT **(a)**, WT + inj *cdk9* mRNA **(b)**, and *mir430*^-/-^ + inj MiR430 **(c)** embryos at 256-cell stage. Shown are representative images of individual nuclei, extracted from a spinning disk confocal microscopy time lapse. The micrographs shown in the main figure are highlighted in green.

**Extended Data Figure 13.**
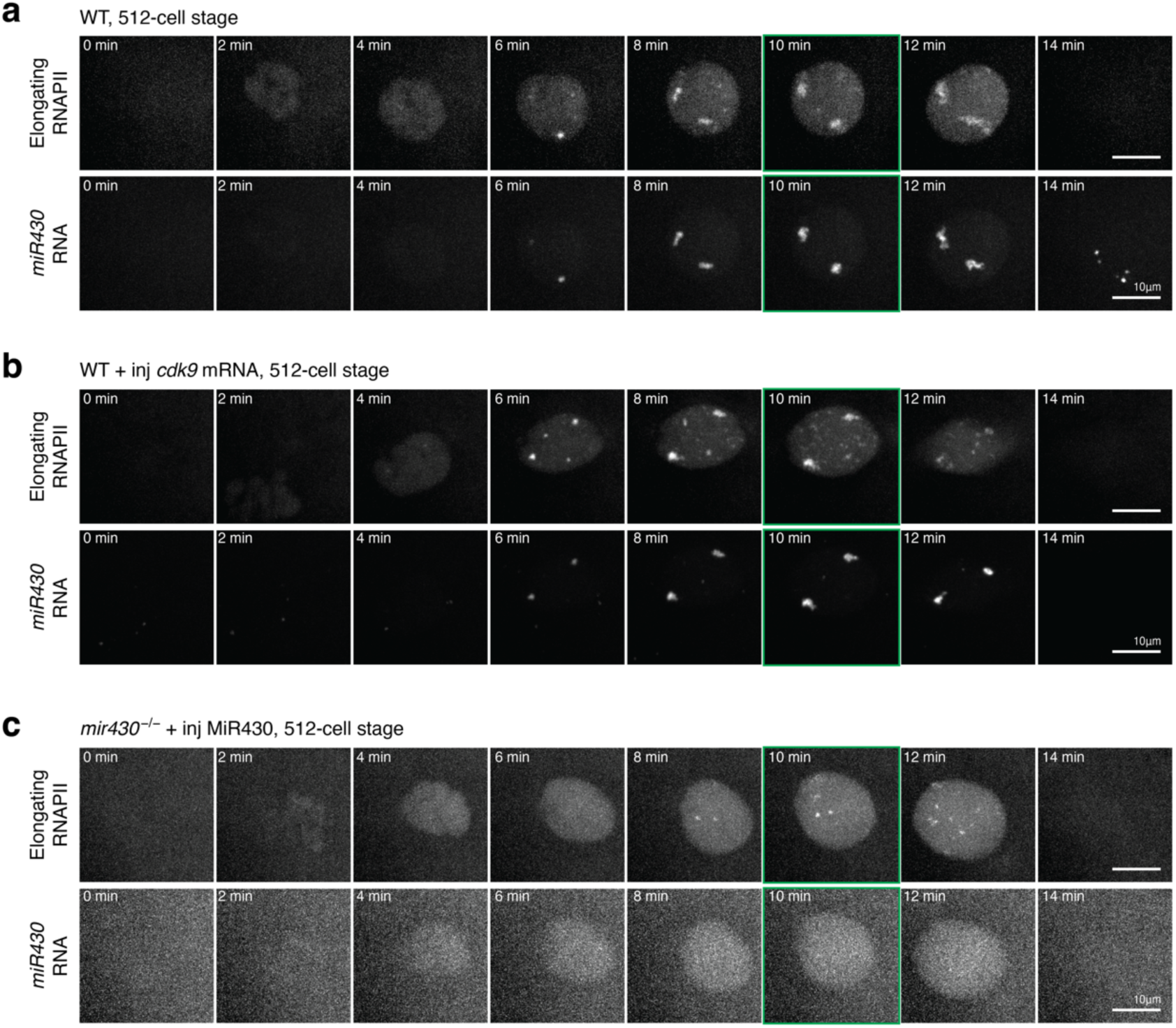
RNAPII-Ser2P signal in WT, WT + inj *cdk9* mRNA and *mir430*^-/-^ + inj MiR430 embryos at 512-cell stage. Visualization of RNAPII Ser2P (with Fabs) and *miR430* RNA (with MOVIE) in WT **(a)**, WT + inj *cdk9* mRNA **(b)**, and *mir430*^-/-^ + inj MiR430 **(c)** embryos at 512-cell stage. Shown are representative images of individual nuclei, extracted from a spinning disk confocal microscopy time lapse. The micrographs shown in the main figure are highlighted in green.

**Extended Data Figure 14.**
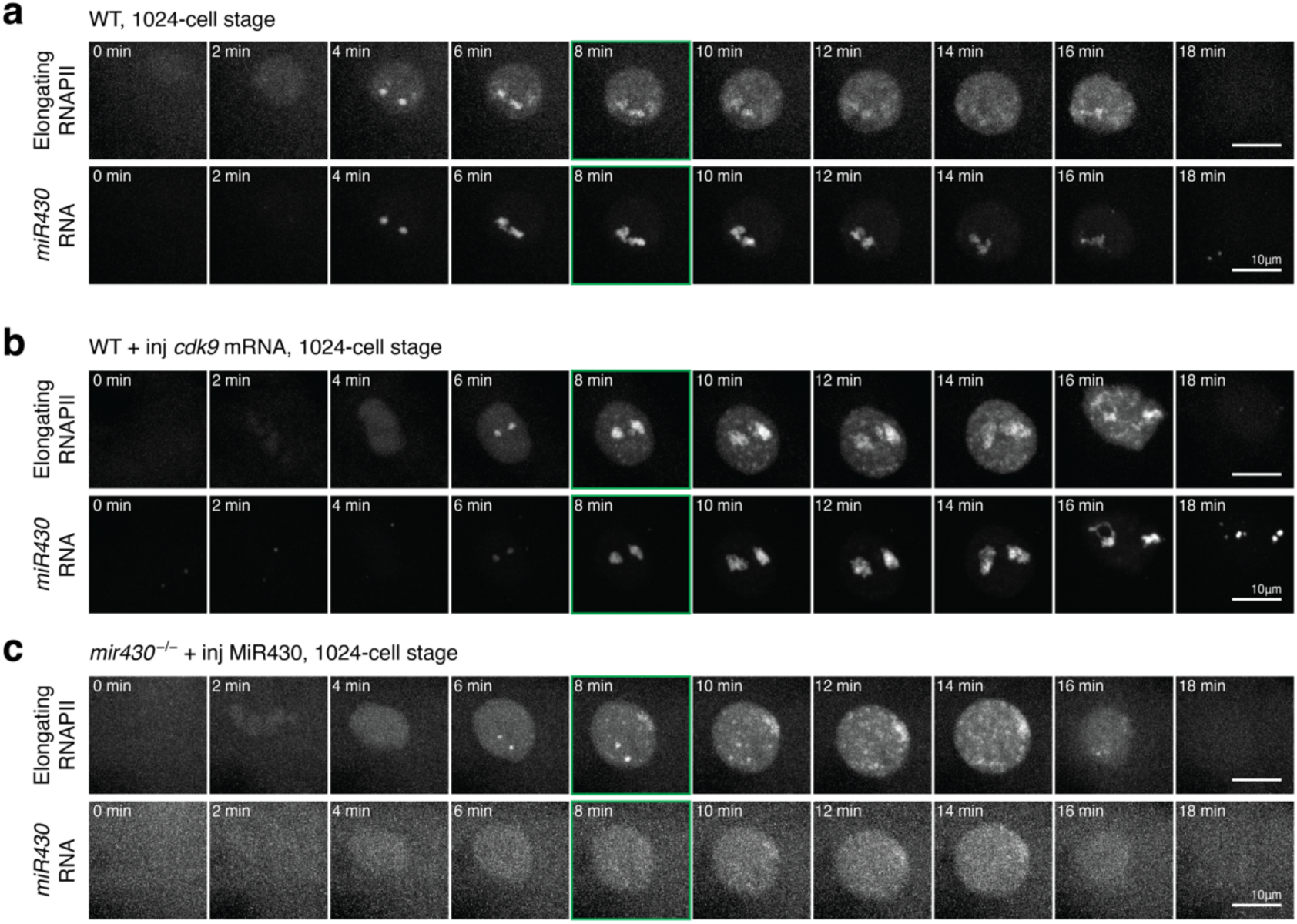
RNAPII-Ser2P signal in WT, WT + inj *cdk9* mRNA and *mir430*^-/-^ + inj MiR430 embryos at 1024-cell stage. Visualization of RNAPII Ser2P (with Fabs) and *miR430* RNA (with MOVIE) in WT **(a)**, WT + inj *cdk9* mRNA **(b)**, and *mir430*^-/-^ + inj MiR430 **(c)** embryos at 1024-cell stage. Shown are representative images of individual nuclei, extracted from a spinning disk confocal microscopy time lapse. The micrographs shown in the main figure are highlighted in green.

**Extended Data Figure 15.**
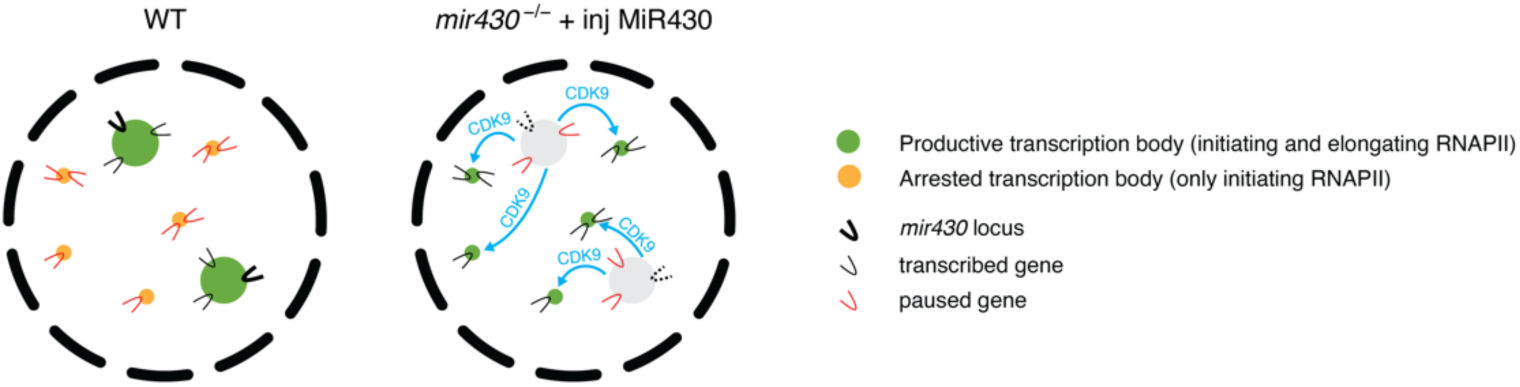
Model for the role of transcription bodies in transcription regulation. In WT nuclei, two large transcription bodies are nucleated by the *mir430* locus. These facilitate the expression of genes that localize there. They also sequester CDK9 (and potentially other factors) which is required for pause release, thereby stalling transcription elsewhere in the nucleus in the initiation state. The specific disruption of the two *mir430* transcription bodies leads to a redistribution of the transcriptional machinery (among which CDK9) which results in the downregulation of genes that localize to the *mir430* transcription bodies and the upregulation of genes elsewhere in the nucleus.

**Extended Data Figure 16.**
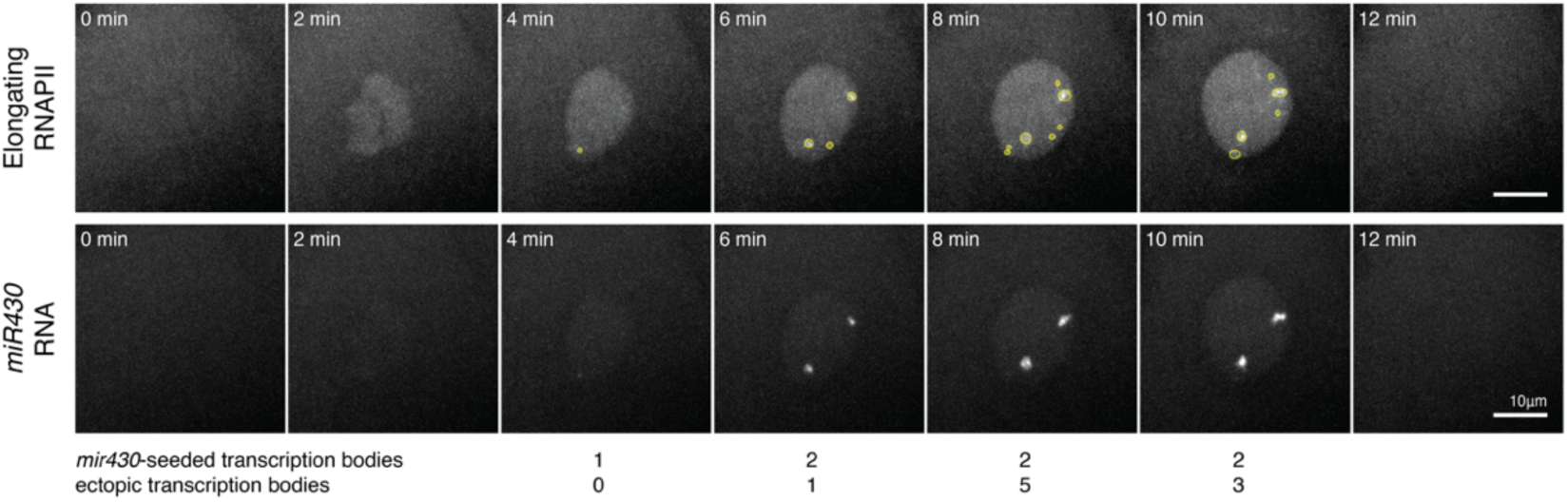
Classification of *mir430* and ectopic transcription bodies. Transcription bodies are identified by detecting elongating RNAPII signal. If this overlaps with MOVIE signal (detecting *miR430* RNA) they are classified as *mir430*, and if they do not overlap with MOVIE signal, they are classified as ectopic transcription bodies. See Methods for more detail.

**Extended Data Table 1.** List of differentially expressed genes in mir430^-/-^ + inj MiR430 *vs.* WT at all stages.

